# Modular organization and selective motifs in the insula provide structural priors for efficient learning

**DOI:** 10.64898/2026.04.22.720260

**Authors:** Songgang Xie, Tao Wang, Ruohan Zhang, Xiaoyu Wang, Ruixinzhu Shao, Xiaofei Wang, Yusi Chen, Henry C. Evrard, Tielin Zhang, Hanfei Deng, Xiong Xiao

## Abstract

The insular cortex (IC) integrates diverse sensory and interoceptive signals to support emotional and cognitive functions, exhibiting topologically functional distinctions. Although long-range pathways of the IC have been extensively mapped, the contribution of intra-IC architecture to network-level information processing remains poorly understood. Here we reconstructed single-cell projectomes of 2,267 IC neurons, generating a comprehensive cellular-resolution map of intra-IC connectivity. Graph-theoretic analyses revealed a hierarchical, topographically organized modular network with a hub-and-spoke organization and selective motif enrichment. Embedding these anatomical priors into recurrent neural network models demonstrated that intrinsic IC topology confers a distinct computational advantage: across multiple cognitive tasks, IC-initialized networks learned faster and exhibited better robustness to perturbations than those initialized with shuffled, randomized, or neighboring somatosensory cortical architectures. Together, these findings identify the IC as a structurally specialized cortical hub whose internal wiring supports efficient learning and provides design principles for brain-inspired networks.

## Introduction

The computational capacity of the brain depends on not only long-range projections linking distant brain regions but also intricate local circuits that shape information processing within each brain region^1–3^. Among various cortical areas, the insular cortex (IC or insula) stands out as a hub bridging interoceptive, limbic, and executive systems, enabling it to integrate external, visceral, and affective signals and transform them into goal-directed actions^4–12^. The IC has been linked to diverse functions, including taste sensation, autonomic regulation, emotion, motivation, timing, and decision making^4,6,8,12,13^. Much of this understanding comes from studies of its long-range efferent or afferent projections^9,10,14–18^. However, how information is organized within the IC, and whether intra-IC connectivity contributes directly to its computational specialization, remain unclear.

Anatomically, the IC is organized along a rostro-caudal axis with systematic variation in cytoarchitecture, gene expression, and projection patterns^4,19–22^. This global IC organization has been linked to its functional specialization across the IC, with more posterior territories preferentially associated with interoceptive and sensory processing^9,16,23,24^, and more anterior territories more strongly implicated in higher-order integration and cognitive control^25–28^. This rostrocaudal functional gradient implies a key role for intra-IC connectivity in linking posterior representations to anterior integrative and decision-related codes, thereby supporting complex behavior^3,29,30^. Indeed, human imaging and lesion studies indicate that functional interactions between posterior and anterior IC underpin awareness, empathy, and adaptive control^7,13,31–33^. Yet, despite this conceptual importance, the precise anatomical organization of intra-IC circuits has remained elusive.

In this study, we employed sparse-labeling methods combined with fluorescence micro-optical sectioning tomography (fMOST)^34,35^ to comprehensively map the connectivity of single IC neurons. We reconstructed the axonal and dendritic morphology of 2,267 individual IC neurons, covering the entire rostrocaudal extent of the IC. This unprecedented dataset allowed us to delineate both intra- and extra-IC connectivity with cellular precision. We uncovered a highly modular intra-IC architecture enriched with small-world-like motifs. By integrating these anatomical data with computational modeling, we show that the intrinsic wiring of the IC provides an optimized architecture for efficient learning on multiple cognitive tasks, illuminating how intra-IC connectivity may support efficient learning.

## Results

### Reconstruction and characterization of single-neuron projectomes of IC neurons

Using established approaches^36,37^, we reconstructed the whole-brain axonal and dendritic morphology of 2,267 sparsely labelled IC neurons at submicrometer resolution using fMOST (Fig. 1a,b). All reconstructed neurons were spatially registered to the Allen Mouse Brain Common Coordinate Framework (CCF), and the IC was parcellated into the anterior (aIC), medial (mIC), and posterior (pIC) IC subregions (Fig. 1c). We computed pairwise morphological dissimilarities using the FNT-dist algorithm^37^ and performed unsupervised hierarchical clustering, yielding 12 distinct projectome-defined neuronal subtypes (Fig. 1d-f). These clusters segregated predominantly into intratelencephalic (IT), pyramidal tract (PT), and corticothalamic (CT) neurons, each displaying characteristic laminar origins and projection topographies (Fig. 1f-h and Extended Data Fig. 1a-k).

**Fig. 1.**
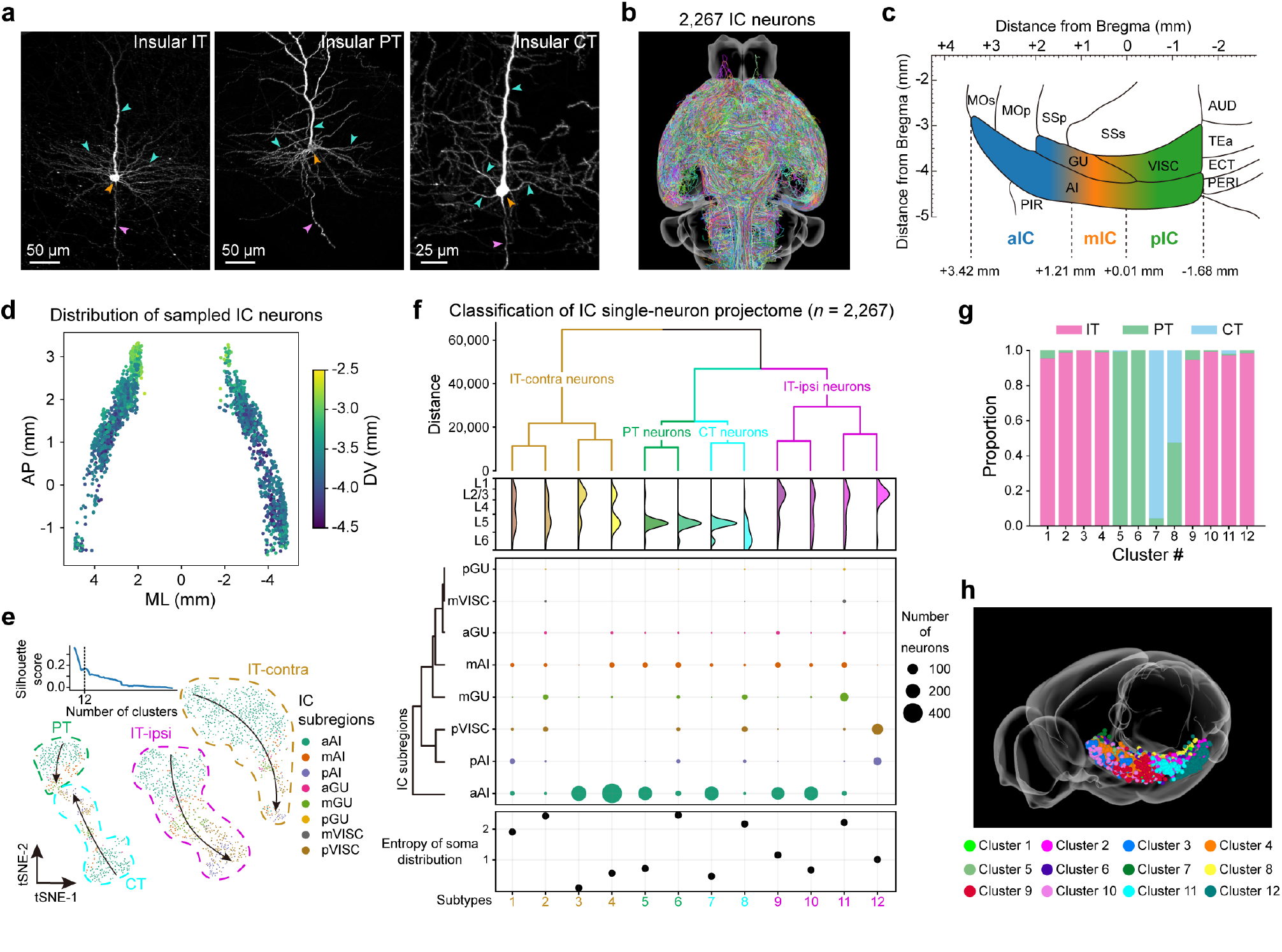
Single-neuron projectome–based classification of insular cortex (IC) neurons. **a**, Representative examples of intratelencephalic (IT), pyramidal tract (PT), and corticothalamic (CT) IC projection neurons. Arrowheads denote soma (orange), dendrites (cyan), and axons (magenta). **b**, Whole-brain reconstructions of 2,267 sparsely labeled IC neurons imaged with fMOST. **c**, Schematic of the IC along the anterior–posterior (AP) axis, delineating anterior (aIC), middle (mIC), and posterior (pIC) subdivisions. **d**, Axial view of soma locations for all reconstructed IC neurons, color-coded by somatic depth (DV). **e**, Top left, silhouette analysis identifying 12 as the optimal number of clusters. Bottom, t-SNE visualization of FNT-dist–based projectome similarity, resolving four major classes—IT-contralateral, IT-ipsilateral, PT, and CT. IT-contralateral class encompasses IT neurons with axonal projections largely targeting the contralateral hemisphere, whereas IT-ipsilateral class consists of neurons projecting almost exclusively to ipsilateral regions relative to their soma. Arrows indicate a global projection gradient aligned with the AP axis. **f**, Hierarchical clustering of projectome similarity across all neurons. Top, dendrogram of the 12 identified subtypes. Middle, distributions of soma locations across cortical layers (upper) and IC subregions (lower). Bottom, entropy of somatic localization across 8 IC subregions, where higher entropy reflects lower regional specificity. **g**, Proportions of IT, PT, and CT neurons within each of the 12 projectome-defined clusters. **h**, Three-dimensional rendering of soma positions for all clusters, with each subtype color-coded distinctly.

IC neurons exhibited pronounced topological organization in both local morphology and collateral architecture along the rostrocaudal axis. Across IT, PT, and CT classes, aIC neurons showed anteriorly tilted dendrites, whereas pIC neurons displayed dendrites aligned more closely with the laminar direction, with mIC neurons showing intermediate configurations (Extended Data Fig. 2a-f). Dendrites and local axons also exhibited projection-class-specific laminar distributions, with IT neurons concentrated mainly in layers 2/3 and 5, PT neurons in layer 5, and CT neurons in layers 5–6 (Extended Data Fig. 2b). First-order collateral analysis identified five IT, three PT, and two CT collateral topologies, consistent with distinct subtype-specific output-distribution strategies (Extended Data Fig. 2g-i). Collateral expansion was greatest in IT neurons and varied further across IC subregions and projectome-defined clusters (Extended Data Fig. 2j,k). Together, these results reveal a rostrocaudally organized anatomical gradient in the IC, in which neuronal position and projection class predict local arbor geometry and collateralization strategy.

### Topographic input-output organization of the IC

Sensory afferents reached the IC through multiple cortical and subcortical pathways, but their terminal distributions were modality-specific and spatially non-uniform. Single-neuron reconstructions revealed modality-specific topographic and laminar organization of visual (VIS), auditory (AUD), olfactory (OLF), somatosensory (SS) cortices, and the parabrachial nucleus (PBN) inputs to the IC (Fig. 2a and Extended Data Fig. 3a-j). VIS inputs were sparse and weak, with limited topographic bias, and preferentially innervated layer 6 of the pIC and layer 2/3 of the aIC (Extended Data Fig. 3a,b). AUD inputs were more prevalent and densely targeted the dorsal pIC, with weaker projections to the aIC; they innervated multiple pIC layers and were enriched in layers 2/3 and 5 in the aIC (Extended Data Fig. 3c,d). OLF inputs formed the strongest cortical afferents, concentrating in the ventral IC and showing a clear topographic gradient from aIC to pIC according to soma position, while preferentially targeting layers 1, 2/3, and 5 and avoiding layer 4 (Extended Data Fig. 3e,f). SS inputs broadly innervated the pIC and dorsal mIC/aIC, also with clear topographic organization, and densely occupied dorsal IC layers, especially in the pIC (Extended Data Fig. 3g,h). Although fewer in number, PBN neurons formed relatively dense focal projections to the AIp and GU and, like OLF inputs, largely avoided layer 4 (Extended Data Fig. 3i,j). Together, these findings indicate that sensory access to the IC follows modality-specific spatial and laminar principles, establishing a topographically organized input architecture across IC subregions.

**Fig. 2.**
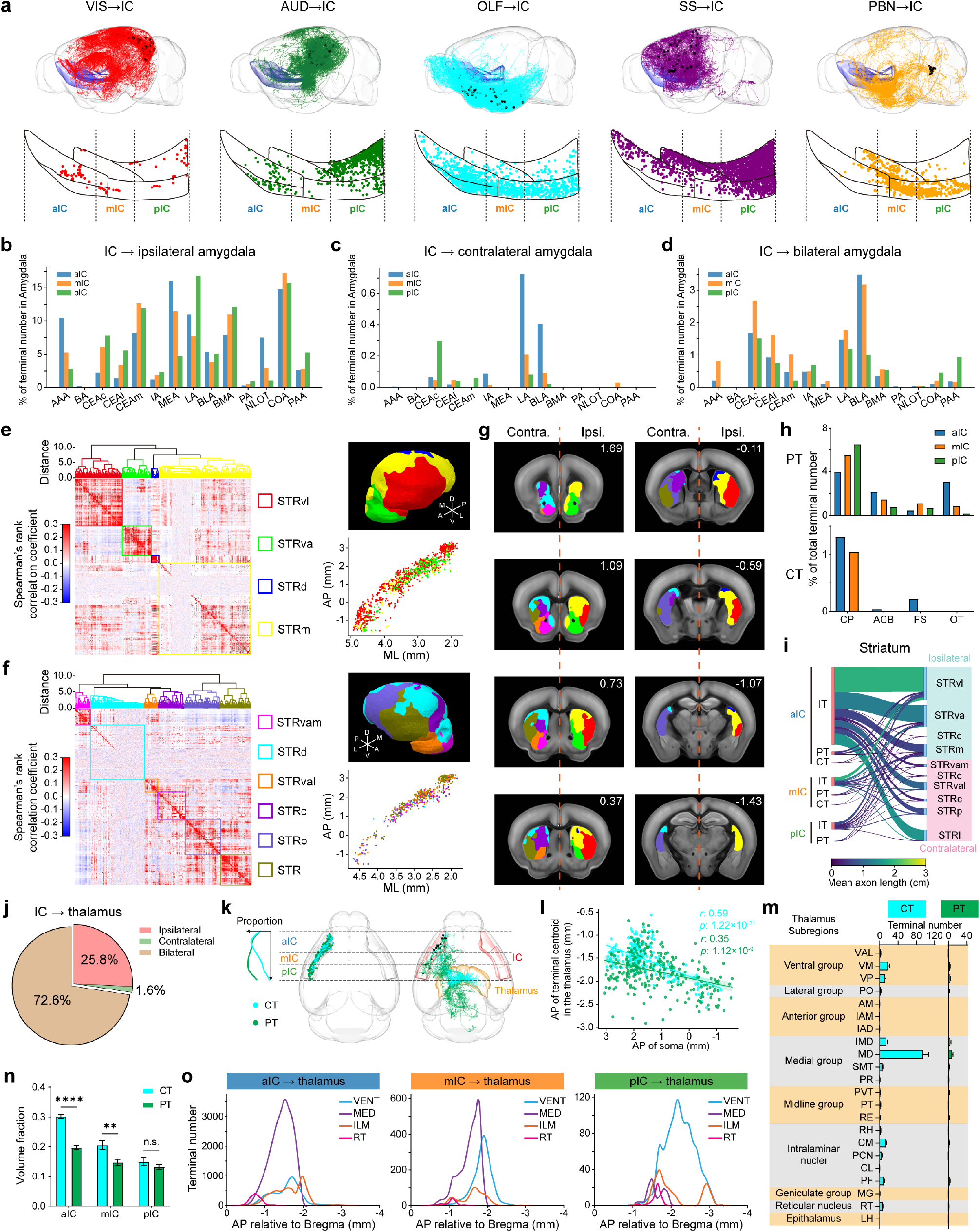
Topographic input-output organization of the IC. **a**, Top, representative VIS, AUD, OLF, SS, and PBN neurons projecting to the IC. Somata are shown in black; IC outlines in blue; axons in red (VIS), green (AUD), cyan (OLF), purple (SS), and orange (PBN). Bottom, topographic maps of VIS-, AUD-, OLF-, SS-, and PBN-derived terminals within the IC. **b-d**, Projection intensities from aIC, mIC, and pIC to ipsilateral (**b**), contralateral (**c**), and bilateral (**d**) amygdalar subregions, expressed as the proportion of terminals within each amygdalar subregion relative to total amygdalar terminals originating from the corresponding IC subdivision. **e**, Left, heatmap of striatal cubes clustered into four subdomains based on ipsilateral projection patterns of IT neurons. Top right, three-dimensional reconstruction of four ipsilateral striatal subdomains. Bottom right, spatial distribution of somata of IC neurons preferentially projecting to each subdomain, color-coded by target domain. **f**, Same as **e** except for contralateral striatal projections. **g**, Eight coronal sections illustrating ipsilateral and contralateral striatal subdomains, with AP coordinates indicated in the upper-right corners. **h**, Projection intensities of PT and CT neurons to four major striatal subregions. **i**, Sankey diagram showing overall projection patterns from IC to bilateral striatal subdomains. Edge width denotes total projection intensity; color scale represents mean projection intensity, operationalized as mean axon length. **j**, Proportion of IC neurons projecting to the ipsilateral, contralateral, or bilateral thalamus. **k**, Left, distribution of the proportions of IC CT and PT neurons projecting to the thalamus along the AP axis. Middle, spatial distribution of somata of thalamus-projecting CT and PT neurons. Right, Projectome of representative thalamus-projecting CT (cyan) and PT (green) neurons (n=10 for both CT and PT neurons). **l**, Linear correlation between AP coordinates of thalamic terminal centroids and somatic AP coordinates of CT/PT neurons. **m**, Projection intensities of CT and PT neurons to thalamic subregions, quantified by terminal counts in each subregion. **n**, Axonal extent within the thalamus for CT and PT neurons, quantified by volume fraction (see Methods; two-way ANOVA followed by Bonferroni’s multiple comparisons; n.s., P > 0.05; **P < 0.01; ****P < 0.0001). **o**, AP-axis distributions of terminals from aIC, mIC, and pIC neurons within the primary thalamic subregions targeted by the IC.

A similarly ordered architecture was evident in IC outputs. Projections from the aIC, mIC, and pIC to downstream limbic and subcortical structures were highly organized and topographic rather than uniform (Fig. 2b-m and Extended Data Fig. 4a-m). Interhemispheric projection mode followed a clear rostrocaudal gradient, with ipsilateral preference increasing from aIC to pIC, bilateral-projecting neurons enriched in the aIC, and these bilateral neurons preferentially located in deep layers (Extended Data Fig. 4a-f). This projection mode also depended on projection class and target, with IT neurons projecting mainly ipsilaterally, CT neurons showing broader bilateral projections to major thalamic targets, and PT neurons exhibiting more heterogeneous patterns across subcortical regions (Extended Data Fig. 4g-i). Within bilaterally projecting IC neurons, distinct clusters differed in soma depth, contralateral terminal depth, and contralateral targeting, and preferentially innervated homotypic contralateral IC subregions, revealing fine-scale organization of interhemispheric IC outputs (Extended Data Fig. 4j-m).

Amygdala-directed outputs showed clear subregion- and laterality-specific biases: ipsilateral projections from all three IC subdivisions converged most strongly onto the lateral and cortical amygdala, but also exhibited distinct preferences across additional amygdalar nuclei, whereas contralateral and bilateral projections were weaker overall and more selectively distributed (Fig. 2b-d and Extended Data Fig. 5a-f). Striatal outputs were likewise organized into discrete topographic channels (Fig. 2e-g and Extended Data Fig. 5g-n). Clustering of IT axon distributions identified four ipsilateral and six contralateral striatal subdomains defined by IC input patterns (Fig. 2e-g). IC neurons exhibited strongly topographically organized projections to the striatum (Extended Data Fig. 5j–l), indicating that distinct IC locations map onto distinct striatal territories. PT and CT neurons projected predominantly to the caudoputamen, with weaker innervation of other striatal subregions (Fig. 2h), and the overall IC-to-striatum output pattern revealed structured routing from specific IC subdivisions and cell classes to defined ipsilateral and contralateral striatal subdomains (Fig. 2i).

Thalamic outputs also exhibited strong organization. Most thalamus-projecting IC neurons showed bilateral projection patterns (Fig. 2j), and these projections were mediated primarily by CT and PT neurons (Fig. 2k). Intra-IC and thalamo-IC projections exhibited distinct laminar preference patterns, supporting separable feedforward- and feedback-like interaction modes between the IC and thalamus (Extended Data Fig. 6a-d). The AP positions of thalamic terminal centroids were significantly correlated with the AP positions of their somata, demonstrating topographic mapping from IC location to thalamic target position (Fig. 2l). CT neurons contributed substantially stronger projections than PT neurons to multiple thalamic subregions, particularly within the ventral, medial, intralaminar, and reticular thalamic groups (Fig. 2m), and occupied larger volumes within the thalamus in the aIC and mIC (Fig. 2n). Moreover, thalamic terminal distributions varied systematically across IC subdivisions, with aIC, mIC, and pIC neurons preferentially targeting distinct AP domains within major thalamic recipient regions (Fig. 2o). Beyond this topography, IC-thalamus connectivity exhibited a clear hierarchical organization, with IC modules differing in their relative hierarchical positions and in how strongly they participated in thalamic feedforward versus feedback interactions (Extended Data Fig. 6e-h).

Together, these analyses identify a coordinated topographic input-output architecture of the IC. Across the IC axis, neuronal position predicts not only how sensory signals enter the IC, but also how information is routed to amygdalar, striatal, and thalamic targets. This organization suggests that the aIC, mIC, and pIC are embedded in distinct yet interconnected anatomical pathways that transform and distribute different classes of information across brain-wide networks.

### Hierarchical modular organization of intra-IC connectivity

Classical models propose that sensory information enters the IC network through the pIC, is transformed across the mIC, and converges within the aIC for integration and behavioral outputs^38–41^. This suggests a hierarchically organized network within the IC. To test this idea at single-neuron resolution, we systematically analyzed the projectomes of neurons with intra-IC projections. We identified two major populations that reciprocally linked the aIC and pIC (Fig. 3a,b and Extended Data Fig. 7a-f). The first, pIC-to-aIC projecting neurons (P2A), were predominantly located in cluster 12 and exhibited strictly ipsilateral projections (Fig. 3a and Extended Data Fig. 7g). The second, aIC-to-pIC projecting neurons (A2P), were enriched in cluster 4 and projected bilaterally (Fig. 2b and Extended Data Fig. 7g). P2A neurons preferentially innervated ipsilateral motor and somatosensory cortices, whereas A2P neurons provided more balanced bilateral projections to motor areas (Extended Data Fig. 7g). These distinct collateralization patterns suggest that P2A and A2P neurons may participate in different information-routing regimes, with P2A neurons biased toward ipsilateral sensorimotor-associated outputs and A2P neurons toward more bilateral motor-associated outputs (Extended Data Fig. 7h). Although both populations lacked clear topographic organization (Extended Data Fig. 7i-l), A2P neurons with more caudal somata consistently produced stronger projections to the pIC (Extended Data Fig. 7k).

**Fig. 3.**
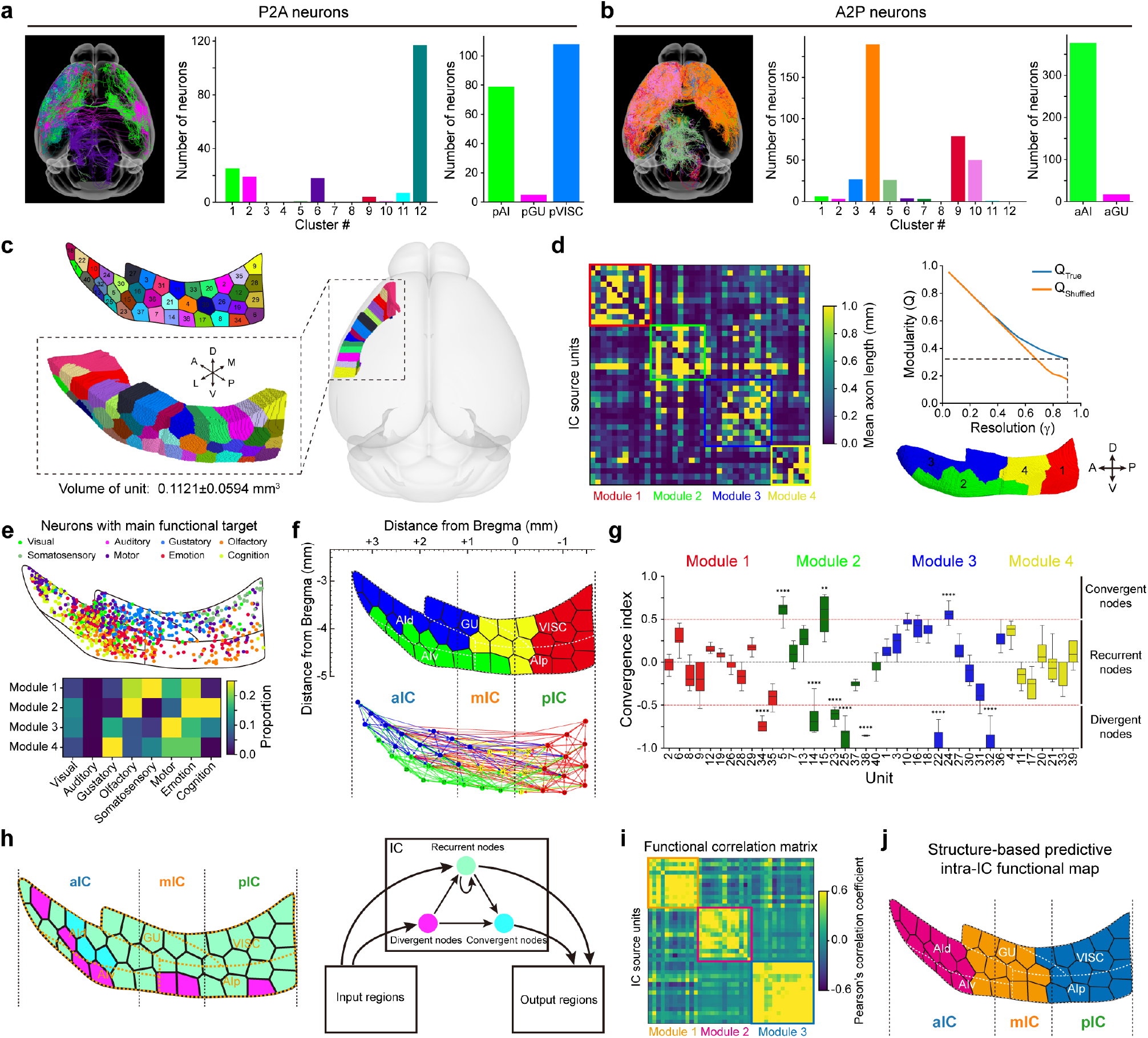
Intra-IC connectivity reveals a modular architecture. **a**, Left, projectome of pIC→aIC (P2A) neurons. Middle, distribution of P2A neurons across the 12 projectome-defined clusters. Right, distribution of P2A somata across pIC subregions. **b**, Same as **a** except for aIC→pIC (A2P) neurons. **c**, Parcellation of the IC into 40 contiguous cortical units with approximately equal surface areas (mean ± s.d. volume: 0.1121 ± 0.0594 mm^3^), each numerically labeled and color-coded. **d**, Left, connectivity heatmap showing clustered projection intensities between IC units. Right, Louvain community detection identifies four structural modules at γ = 0.9, where modularity (Q) of the true matrix (blue) maximally exceeds the shuffled control (orange). Data are presented as mean ± s.e.m. Bottom right, sagittal view of the four modules. **e**, Top, single-neuron functional topography of cortical projection targets (visual, auditory, gustatory, olfactory, somatosensory, motor, emotion, cognition); neurons downsampled 0.5×. Bottom, module-specific functional preferences. **f**, Top, spatial layout of the four modules overlaid on Allen Mouse Brain Atlas subdivisions. Bottom, intra-IC network graph showing connections with projection intensity ≥ 0.4. **g**, Convergence index distributions for all units, grouped by module. Red dashed lines mark thresholds for convergent (≥ 0.5), recurrent (−0.5 to 0.5), and divergent (≤ −0.5) nodes. **P < 0.01; ****P < 0.0001; one-sided t-test against ≥ 0.5 or ≤ −0.5. **h**, Left, spatial distribution of divergent, recurrent, and convergent nodes across the IC; orange dashed boundaries denote the five IC subregions. Right, schematic model of proposed intra-IC processing pathways linking input, recurrent, and output nodes. **i**, Pearson correlation matrix of unit-wise activity derived from the rate-based whole-IC model, revealing three functional modules (γ = 0.75). **j**, Structure-based functional map predicted from intra-IC connectivity, delineating three regions corresponding to the functional modules in **i** and aligning with classical aIC, mIC, and pIC compartments.

To achieve a fine-grained analysis of intra-IC connectivity, we first parcellated the entire IC surface into 40 spatially contiguous subregions of approximately equal area using K-means clustering^36,37^. This surface-based parcellation was subsequently volumetrically extended to encompass the full IC, yielding 40 discrete units for connection analysis (Fig. 3c). We quantified projection intensities between all unit pairs, followed by community detection using the Louvain algorithm^37^ (see Methods). We identified four major structural modules that closely corresponded to the dorsal and ventral agranular IC (AId and AIv), mIC, and pIC (Fig. 3d). To infer potential functional specializations, we quantified projection intensities from each structural module to functionally defined cortical domains. This analysis revealed distinct associations: module 1 was primarily linked to olfactory, somatosensory, and emotional networks; module 2 to olfactory, emotional, and cognitive regions; module 3 to motor-related areas; and module 4 to gustatory circuits (Fig. 3e).

Inter-unit connectivity exhibited substantial heterogeneity in the balance of afferent inputs and efferent outputs across IC units. To quantify these connectivity asymmetries, we computed a convergence index for each unit, classifying nodes with values > 0.5 as convergent, < –0.5 as divergent, and intermediate values as recurrent (see Methods). Divergent nodes were broadly distributed across the aIC, mIC, and pIC with a ventral bias, whereas convergent nodes were confined to the aIC (Fig. 3f,g). Integrating these findings with classical anatomical models, we propose a revised framework for intra-IC connectivity (Fig. 3h). Divergent nodes may act as broadcasting nodes that distribute incoming signals across multiple targets for parallel processing. Recurrent nodes may serve as integrative centers, maintaining feedback loops within the network while relaying secondary outputs to downstream regions. Convergent nodes, situated in the aIC, are positioned to integrate information arriving from divergent and recurrent nodes to generate strong, focused outputs to extra-IC targets (Fig. 3h).

To test whether the static intra-IC connectivity is sufficient, by itself, to give rise to functional modularity of neural activities, we constructed a rate-based network model^42^ based on the reconstructed intra-IC connection matrix (Extended Data Fig. 7m) and its inter-unit distance information (Extended Data Fig. 7n; see Methods). Simulated current inputs were supplied to the neural network to mimic external sensory drive. The functional connectivity matrix (Fig. 3i) was then generated as pairwise Pearson correlations of simulated neural activities. Community detection of this matrix using the Louvain algorithm revealed three functionally distinct modules that closely corresponded to the anatomically defined aIC, mIC, and pIC subregions (Fig. 3j). These simulations indicate that the reconstructed intra-IC connectivity is sufficient to generate activity modules aligned with major IC subdivisions, consistent with a hierarchically modular organization of the IC.

### IC architecture accelerates learning in RNN

Extensive evidence implicates the IC in the process of action selection and decision-making^9,15,43,44^. Guided by the principle that anatomical structure can constrain computation^45,46^, we next examined whether the intrinsic architecture of the IC provides structural priors that facilitate rapid learning. We constructed recurrent neural networks (RNNs) with different initializations trained on a go/no-go task^47^ (Fig. 4a; see Methods). Pre-generated Poisson spike trains served as sensory inputs delivered to the network. During training, only the input weights, recurrent weights, and biases of the recurrent layer were updated by gradient descent. To isolate the contribution of initial network topology, we compared four RNN groups that differed only in their initial recurrent connectivity: (1) Insula-origin, empirically derived intra-IC connectivity with additive variability; (2) Insula-shuffle, degree-preserving shuffling of intra-IC connection weights; (3) Random-Fro, absolute-valued Kaiming-normal initialization scaled to match the Frobenius norm of (1); (4) Random-SR, as (3), but scaled to have a matched spectral radius (Extended Data Fig. 8a,b). Ten independent training environments were generated, with each environment containing twenty matched runs per RNN group sharing identical input/output and bias initializations. Although loss trajectories were similar across all groups (Extended Data Fig. 8c), the Insula-origin networks showed an earlier rise in training accuracy (Fig. 4b). Insula-origin networks reached the learning boundary significantly earlier than the other three groups and exhibited a substantially larger area under the curve (AUC) of accuracy throughout training (Fig. 4c), indicating accelerated acquisition of task-relevant decision boundaries. To visualize how network weights evolved over training, we projected recurrent weight matrices into t-SNE space (Fig. 4d). The Insula-origin networks displayed organized early-phase trajectories characterized by initial convergence followed by divergence, a structured pattern absent in the shuffled or random groups (Fig. 4d). Consistently, early-phase weight evolution in the Insula-origin group showed significantly higher clusterability (Fig. 4e), indicating that the biological IC topology induces coordinated and reproducible optimization dynamics.

**Fig. 4.**
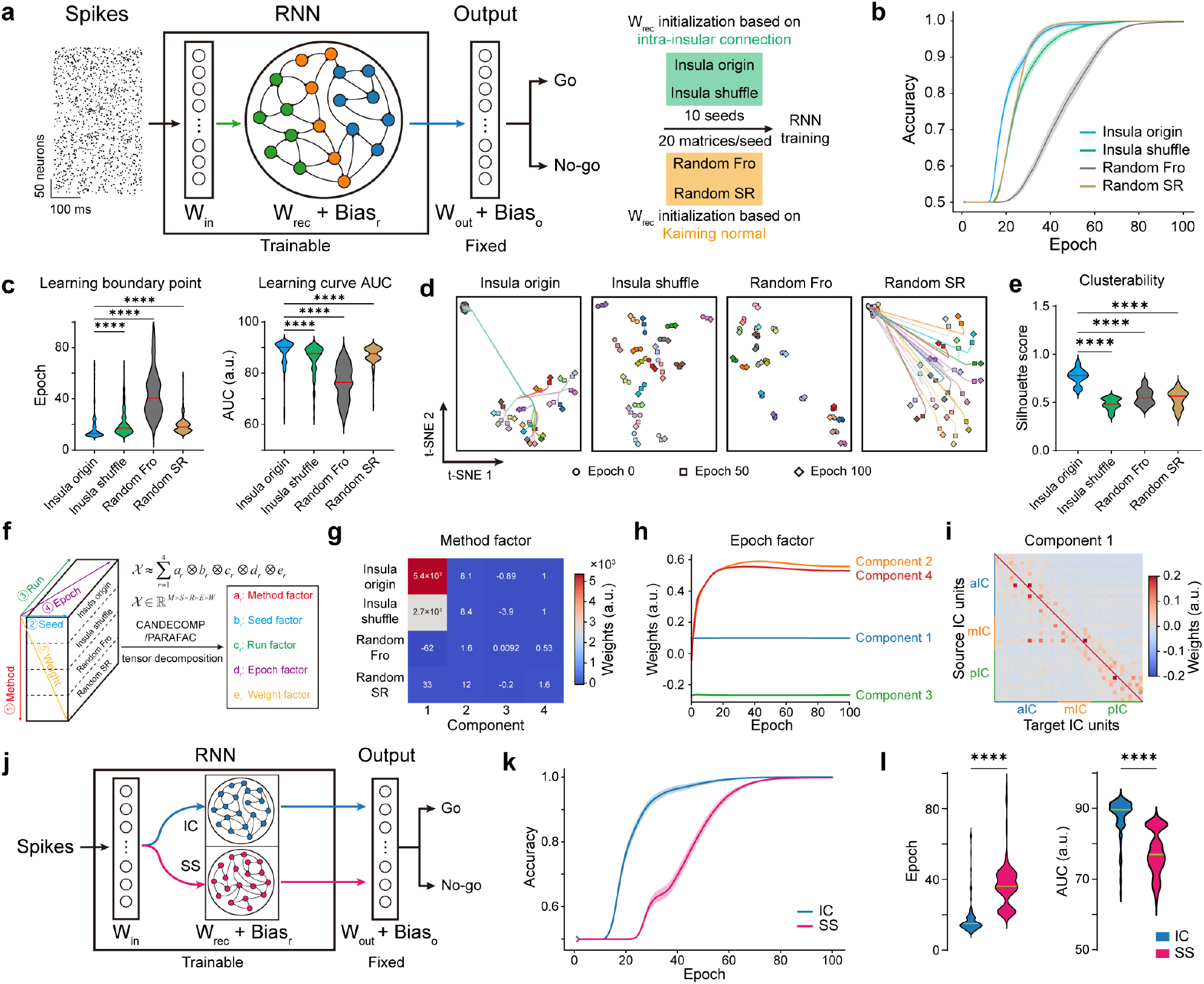
Structural priors in the IC accelerate recurrent network learning. **a**, Schematic of the RNN architecture and training protocol. **b**, Accuracy trajectories of convergent runs across the four RNN groups; Insula-shuffle and Random-SR groups show highly similar optimization profiles. **c**, Left, epochs required to reach the learning boundary (defined as the earliest sustained accuracy increase above 0.5), with lower values indicating faster learning (****P < 0.0001). Right, accuracy AUC for convergent runs (****P < 0.0001). Kruskal-Wallis test followed by Dunn’s multiple comparisons. **d**, Representative t-SNE embeddings of recurrent weight matrices across training for the four groups. Circles, squares, and diamonds denote the states at initialization, epoch 50, and training completion, respectively. **e**, Clusterability of recurrent-weight trajectories during the first 18 epochs; higher values indicate more coherent early-phase weight evolution (****P < 0.0001, one-way ANOVA followed by Dunnett’s multiple comparisons). **f**, Schematic of fifth-order recurrent-weight tensor construction and subsequent low-rank tensor decomposition. **g**, Method-factor loadings across four components for each initialization class. **h**, Temporal evolution of epoch-factor loadings across components over training. **i**, Weight-factor loadings for component 1, revealing the dominant learning strategy selectively engaged by IC-architected RNNs. **j**, Schematic of IC-based versus SS-based architectures used for RNN training. **k**, Accuracy trajectories of convergent runs for IC and SS RNN groups. **l**, Left, epochs required to reach the learning boundary for IC versus SS groups (****P < 0.0001, Wilcoxon rank sum). Right, AUC of accuracy curves for IC and SS groups (****P < 0.0001, Wilcoxon rank sum).

To determine whether the observed acceleration could be explained simply by favorable local curvature properties of the loss landscape, we quantified the spectral radius and the condition number of Hessian matrix during the early-training phase, two key indicators of convergence speed and optimization stability. The spectral radius was comparable among the Insula-origin, Insula-shuffle, and Random-SR groups, yet was markedly lower than the Random-Fro group. In contrast, the condition number showed no significant differences across the four groups (Extended Data Fig. 8d,e). Loss-landscape visualizations further revealed that Insula-origin, Insula-shuffle, and Random-SR groups shared similarly smooth and topographically aligned landscapes (Extended Data Fig. 8f). Thus, generic matrix-level properties cannot account for the superior learning rates of the Insula-origin networks. These results indicate that the faster learning observed in IC-initialized networks depends on specific topological features of the IC-derived connectivity rather than on simple differences in optimization landscape geometry.

To further dissect how this topology shapes learning, we constructed a fifth-order tensor of all recurrent weight matrices and performed low-rank tensor decomposition, yielding five latent factors (Fig. 4f and Extended Data Fig. 8g). Component 1 of the method factor cleanly separated IC-architectured networks (Insula-origin) from shuffled and randomized controls, showing strongly positive loadings in the Insula-origin group, moderate loadings in the Insula-shuffle group, and weak loadings in the others (Fig. 4g). The epoch factor revealed that component 1 and 3 represented the static initialization bias, whereas components 2 and 4 captured training-dependent dynamics (Fig. 4h). These results indicate that IC-based RNNs strongly and consistently adopted a shared, topology-dependent learning strategy encoded in the weight factor, whereas control networks either failed to engage or only weakly engaged this strategy. Component 1 of the weight factor captured the main divergence in learning strategy: IC-architectured RNNs concentrated recurrent-weight updates among spatially proximal nodes, substantially enhancing local information-routing efficiency. Most notably, they selectively strengthened the projection from a key mIC node to the aIC, strengthening this pathway as a dominant relay for downstream information flow (Fig. 4i). In contrast, components 2–4 of weight factor reflected learning strategies shared across all architectures, yet the method factor indicated that their contribution to the overall learning strategy of the RNNs was minimal. Notably, while component 3 shared similarities with component 1 in terms of weight factor, it substantially strengthened connections between adjacent nodes while simultaneously weakening many long-range connections, which would likely impair overall information transmission (Fig. 4g and Extended Data Fig. 8j). Seed factor analysis showed that, with exception of component 2, the learning strategies captured in the weight factor were robust to random initialization (Extended Data Fig. 8h), while run factors confirmed minimal variability across training runs (Extended Data Fig. 8i). Together, these results suggest that IC-derived topology biases learning toward more coordinated and spatially structured recurrent-weight updates.

Because go/no-go task acquisition also involves updates to input weights and biases of recurrent layer, we next evaluated their relative contributions. Rollback tests, in which specific trained parameters were reset to their initial values, revealed that recurrent weights and biases exerted the largest influence on accuracy, hinting that biases may modestly affect learning speed (Extended Data Fig. 8k). Low-rank tensor decomposition of input weights, however, showed no architectural segregation and static epoch factors, indicating negligible contribution to the accelerated learning (Extended Data Fig. 8l,m). By contrast, bias tensor decomposition exhibited weak architectural discrimination and dynamic epoch factors (Extended Data Fig. 8n,o). Nevertheless, direct comparison of early-training dynamics demonstrated that recurrent weights underwent substantially larger updates than biases (Extended Data Fig. 8p). These analyses collectively indicate that recurrent-weight optimization, shaped by the insular topology, is the primary driver of accelerated learning, with biases contributing modestly and input weights contributing minimally.

Next, we asked whether accelerated learning represents a general property of cortical structural priors. Using the same analytical pipeline, we extracted single-neuron projectome (n = 3943 neurons)–derived connectivity from the neighboring somatosensory cortex (SS) (Extended Data Fig. 9a), a neighboring primary sensory region with a distinct anatomical and functional profile from the IC, and used these SS-derived structural priors to initialize new RNNs (Fig. 4j). In contrast to IC-architectured networks, SS-architectured RNNs learned significantly more slowly and exhibited higher training loss (Fig. 4k,l and Extended Data Fig. 9b). Together, these results indicate that faster learning is not a universal consequence of cortical structural priors, but instead depends on specific topological features of IC-derived connectivity.

### Task-dependent benefits of IC architecture in RNN learning

We next asked whether the learning advantage conferred by IC-derived topology extended beyond the go/no-go task to task domains with distinct computational demands. To address this, we evaluated four RNN groups on five representative tasks: delayed match-to-sample, interval reproduction, context-dependent sensorimotor mapping, four-armed bandit reinforcement learning, and Penn Treebank language modeling (Fig. 5a-r). The benefits of IC architecture proved to be task-dependent rather than universal. In the delayed match-to-sample task, RNNs initialized with the IC architecture showed faster loss reduction and higher task accuracy than all control groups, indicating a clear advantage for maintaining and comparing short-term stimulus representations (Fig. 5a-d). A similar advantage was observed in the interval reproduction task, in which IC-initialized RNNs achieved lower loss and smaller timing error than control networks, consistent with improved short-term temporal maintenance and reproduction (Fig. 5e-g).

**Fig. 5.**
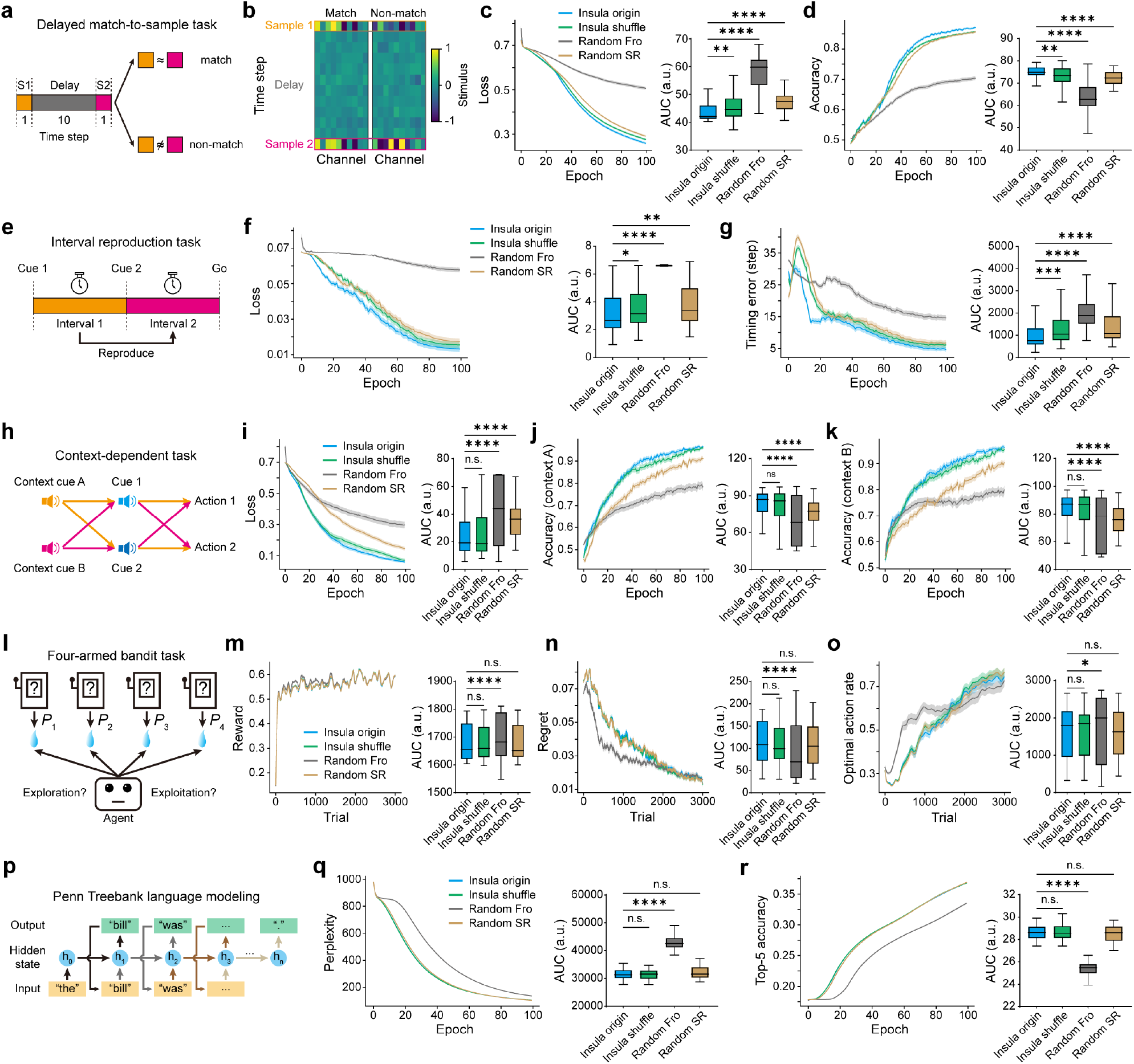
Task-dependent benefits of IC architecture in RNN learning. **a**, Schematic of the delayed match-to-sample (DMS) task. S1 and S2 represent the first and second sample stimuli, respectively. Trials are defined as match (S1 and S2 are similar) or non-match (S1 and S2 are dissimilar). **b**, Input examples for match and non-match trials. **c**,**d** Left, training loss (**c**) and accuracy (**d**) for the four RNN groups in the DMS task. Right, corresponding statistical comparison of the AUC across the four groups. **e**, Schematic of the interval reproduction task. The agent is required to memorize the time interval between cue 1 and cue 2 and then reproduce this interval after cue 2 ends. **f**,**g** Left, loss (**f**) and timing error between the reproduced and actual intervals (**g**) of the four RNN groups in the interval reproduction task. Right, corresponding statistical comparison of the AUC across the four groups. **h**, Schematic of the context-dependent task. In context A, cue 1 corresponds to action 1 and cue 2 to action 2; in context B, cue 1 corresponds to action 2 and cue 2 to action 1. **i-k**, Left, loss (**i**), accuracy under context A (**j**), and accuracy under context B (**k**) of the four RNN groups in the context-dependent task. Right, corresponding statistical comparison of the AUC across the four groups. **l**, Schematic of the four-armed bandit task. The environment contains four slot machines, each with a distinct fixed reward probability. On each trial, the agent interacts with one machine to attempt to obtain a reward, with the goal of maximizing total rewards within limited trials. **m-o**, Left, average reward (**m**), regret (**n**), and optimal action rate (**o**) of the four RNN groups in the four-armed bandit task. Right, corresponding statistical comparison of the AUC across the four groups. **p**, Schematic of the RNN-based self-supervised language modeling task on the Penn Treebank (PTB) dataset. **q**,**r** Left, perplexity (**q**) and top-5 accuracy (**r**) across training for the four RNN groups in the PTB language modeling task. Right, corresponding statistical comparison of the AUC across the four groups. Data are presented as mean ± s.e.m. n.s., P>0.05; *, P<0.05; **, P<0.01; ***, P<0.001; ****, P<0.0001. Kruskal-Wallis test followed by Dunn’s multiple comparisons.

In the context-dependent task, IC-initialized RNNs also outperformed the two random-control groups, but performed similarly to the insula-shuffled networks (Fig. 5h-k). This result suggests that, for this task, performance depends less on precise higher-order IC topology and more on lower-order structural statistics preserved by shuffling. By contrast, the IC architecture did not confer a clear advantage in the four-armed bandit task or the Penn Treebank language modeling task, in which IC-based and shuffled networks performed similarly to one another and only modestly differed from random controls (Fig. 5l-r). Thus, the computational benefit of IC-derived topology was selective rather than universal, emerging most strongly in tasks that require rapid feature extraction, short-term maintenance, and comparison, but not in tasks dominated by long-range credit assignment or sequential dependency learning.

### IC architecture exhibits a hub-and-spoke modular organization with selective motif enrichment

The superior learning performance of IC-initialized RNNs relative to shuffled, randomized, and SS networks suggested that the IC topology contains distinctive structural features. To identify them, we converted recurrent weight matrices from RNN groups into directed graphs and extracted their topological features using a directed graph convolutional network (DGCN)^48^ (Fig. 6a). When projected into t-SNE space, the resulting embeddings showed a clear separation of the Insula-origin group from the remaining three groups (Fig. 6b). However, a linear logistic-regression classifier trained on the DGCN embeddings failed to distinguish between Insula-origin networks and Insula-shuffle networks (Fig. 6c). These results indicate that the IC network contains topological structure not captured by random networks, yet shares some topological properties such as the degree sequence with degree-preserving shuffled networks. Furthermore, using the same method to compare IC, SS, and random networks, we observed distinct differences in global topological features among them (Extended Data Fig. 9c). This indicates that different cortical regions can exhibit substantially different topological architectures. Spectral density analysis of the directed normalized Laplacian further revealed asymmetric eigenvalue distributions in both IC and SS networks (Extended Data Fig. 9d), indicating that biological brain networks are not homogeneous but instead possess distinctive structural features. Compared to random networks, the IC network exhibited a higher proportion of low eigenvalues (Fig. 6d,e and Extended Data Fig. 9d), reflecting stronger community structure and modular organization. Compared to SS network, the eigenvalues of the Laplacian spectrum of IC network are more concentrated around 1.0. This indicates that, while both networks exhibit modular organization, IC network has stronger inter-module connections, leading to better cross-module information transmission and integration. In contrast, SS network shows stronger segregation between modules, which may reflect more specialized functional differentiation. Furthermore, shuffling the IC network resulted in a more uniform and symmetric spectral distribution, indicating disruption of its inherent modular architecture (Fig. 6d,e).

**Fig. 6.**
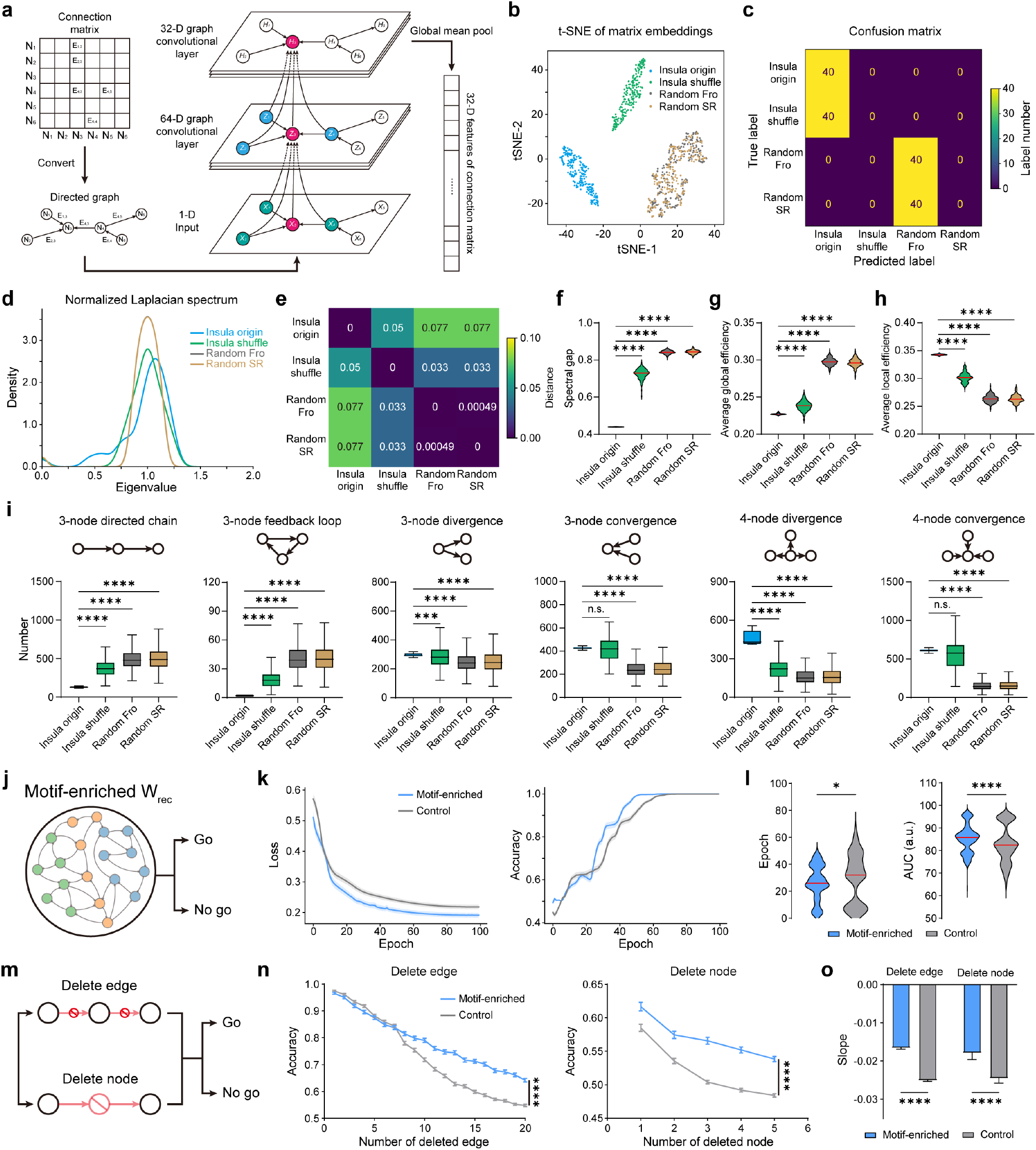
Selective motifs underlie the computational advantage of IC architecture. **a**, Schematic of the directed graph convolutional network (DGCN) used to extract global topological features from recurrent weight matrices. **b**, t-SNE visualization of 32-dimensional DGCN embeddings of initialized recurrent weight matrices from the four RNN types; each point represents one matrix and is color-coded by group. **c**, Confusion matrix of a logistic-regression classifier trained on DGCN embeddings. **d**, Spectral density distributions of normalized Laplacian eigenvalues for the four matrix types. **e**, Pairwise Wasserstein distances between normalized Laplacian spectra of the four groups. **f-h**, Comparisons of spectral gap (**f**), average global efficiency (**g**), and average local efficiency (**h**) across groups (****P < 0.0001, Kruskal-Wallis test followed by Dunn’s multiple comparisons). **i**, Frequencies of six representative 3-node and 4-node motifs across four matrix types (****P < 0.0001, Kruskal-Wallis test followed by Dunn’s multiple comparisons). **j**, Schematic of RNN training using recurrent weights enriched for directionally specific motifs. **k**, Loss and accuracy trajectories for motif-enriched versus control networks during training. **l**, Comparisons of learning boundary (left; ****P < 0.0001) and AUC (right; ****P < 0.0001) between motif-enriched and control networks (Wilcoxon rank sum test). **m**, Schematic of structural perturbations applied to trained networks—edge deletion or node deletion. **n**, Test accuracy under progressively increasing numbers of deleted edges (left; ****P < 0.0001) or deleted nodes (right; ****P < 0.0001). Two-way ANOVA. **o**, Slope of accuracy-decline curves from **n**, showing that motif-enriched networks degrade significantly more slowly than controls (****P < 0.0001, two-way ANOVA followed by Bonferroni’s multiple comparisons).

We next evaluated classical graph-theoretic metrics. Weighted degree distributions in the IC network showed a marked leftward shift and heavy-tailed profile, highly similar to those of the shuffled networks, but distinct from the near-Gaussian distributions of the random networks (Extended Data Fig. 10a-c). This profile indicates that most IC nodes maintain sparse connectivity while a minority function as high-degree hubs, consistent with the hierarchical structure. Compared to shuffled and random networks, the IC network exhibited a smaller spectral gap, lower average global efficiency, longer average shortest path length, higher average local efficiency, and consistently negative assortativity (Fig. 6f-h and Extended Data Fig. 10d,e). These characteristics are consistent with a highly modular organization of the IC network. When compared to the SS network, the IC network demonstrates a higher average clustering coefficient, lower average local efficiency, and lower assortativity, consistent with stronger segregation of local processing alongside preserved large-scale integration (Extended Data Fig. 9e-g). Together, these features indicate that IC architecture combines modular organization, small-world features, and disassortative mixing—properties reminiscent of large-scale human brain networks^49,50^. Applying the PageRank algorithm to the IC network identified three dominant hubs (Extended Data Fig. 10f). These observations are consistent with a hub-and-spoke organizational principle^51,52^, wherein information is locally integrated within modules and selectively routed through specialized hubs that enable rapid inter-module communication—balancing local processing with efficient global integration (Extended Data Fig. 10g).

Finally, motif analysis revealed enrichment of convergent, divergent, and feedforward motifs, coupled with depletion of linear and cyclic chain motifs in the IC network (Fig. 6i and Extended Data Fig. 10h). By comparing the top 10 most enriched motifs in the IC and SS networks, we observed that the IC network predominantly exhibits motifs characterized by unidirectional information flow between nodes, whereas the SS network shows a preference for motifs associated with bidirectional information transfer. This suggests that, at the motif level, the IC contains a stronger bias toward unidirectional information flow than the SS network (Extended Data Fig. 9h-k). Furthermore, we found that selectively enriching specific motifs that are overrepresented in the IC network partially reproduced the motif enrichment pattern of the IC network in the constructed networks. The highest similarity was achieved when enriching the 4-node multi-layer feedforward (4mlf) motif (Extended Data Fig. 10i-l). When networks generated by embedding the 4mlf motif were used as the initial recurrent weights for RNNs trained on a go/no-go task, these networks accelerated learning compared to randomly initialized control networks (Fig. 6j-l). More notably, after structural perturbations were applied to the trained recurrent weights, the motif-enriched group exhibited significantly stronger resilience than the control group (Fig. 6m-o). Together, these results indicate that selective motif enrichment, particularly of multi-layer feedforward motifs, is sufficient to recapitulate part of the computational advantage associated with IC-derived topology.

## Discussion

Here we have established the first single-cell–resolution map of intra-IC connectivity, revealing a highly modular, hierarchically organized network that bridges sensory, interoceptive, and cognitive domains. Although IC circuitry has been studied primarily through its long-range inputs and outputs, our data highlight that the IC is also supported by dense, highly structured internal wiring (Fig. 3d-j and Extended Data Fig. 7a-h). This reciprocal microcircuitry enables information to be perceived, integrated, and transformed within the IC before being transmitted to downstream structures. Such an architecture positions the IC as an internal integrative substrate within the cortex, an anatomical substrate for rapid cross-modal exchange and reconfiguration of ongoing computations according to changing environmental or physiological demands^53^.

The IC’s internal wiring is neither random nor uniform but organized into discrete modules connected through selective convergent, divergent, and feedforward motifs (Fig. 3d, 6i, and Extended Data Fig. 10h). These motifs support hierarchical information flow from posterior to anterior compartments, paralleling a progressive transformation from sensory representation to cognitive inference^3,7,29,30^. The convergence of these features, including small-world topology, modular hierarchy, and motif enrichment, provides a plausible structural basis for flexible and efficient information routing^54,55^. By embedding this architecture into artificial neural networks, we demonstrate that the IC topology itself confers computational advantages: RNNs initialized with IC connectivity learned a decision task faster than equivalently sized shuffled or randomized networks (Fig. 4b-e). This acceleration was not explained by metrics reflecting optimization landscape geometry, such as spectral radius or condition number of Hessian matrix, but emerged from topological structure (Extended Data Fig. 8d-f). Tensor decomposition analyses revealed that IC networks preferentially strengthened short-range connections and established a dominant information highway from the middle to anterior nodes, forming a dynamic “superhub” that optimized local integration and long-range transfer (Fig. 4f-i). These results suggest that the small-world, hub-and-spoke design of the IC supports efficient biological computation, an evolutionary solution that balances wiring cost and efficiency.

From an evolutionary perspective, the emergence of such motifs (Fig. 6i and Extended Data Fig. 10h) may reflect adaptive pressure for circuits capable of rapid learning in multimodal and unpredictable environments^56,57^. The IC integrates visceral, gustatory, somatosensory, and affective signals, domains that demand flexible updating rather than stable memorization. Its architecture may therefore reflect a cortical strategy optimized not for maximal representational capacity, but for adaptive learning. Comparable structural principles may underlie other regions supporting behavioral flexibility, such as the prefrontal cortex^58^, suggesting a conserved blueprint for learning-efficient architectures across the mammalian brain. At the systems level, the IC’s hub-and-spoke organization (Extended Data Fig. 10f,g) implies that functional specialization and integration are dynamically balanced through a small number of connector nodes^51,52^. Perturbations of these hub-like nodes could therefore be expected to influence information flow broadly across the IC network, providing a mechanistic explanation for the IC’s vulnerability in neuropsychiatric disorders characterized by impaired adaptation, such as depression, addiction, and anxiety^4,6,59^. In this light, network measures such as hub centrality or motif balance may provide candidate biomarkers of adaptive potential, and targeted interventions aimed at restoring intra-IC connectivity could improve learning and behavioral flexibility. Together, these findings support the view that the IC contains an internal circuit architecture well-suited for integrating and routing multimodal information, and that this architecture can endow recurrent networks with selective learning advantages. The principles uncovered here, including modular organization, motif-based routing, and topology-dependent learning, may extend beyond the IC, offering a unifying framework for understanding how evolution shaped cortical networks to embed intelligence into their very architecture.

## LIST OF ABBREVIATIONS

AAA: anterior amygdalar area
ACA: anterior cingulate area
ACB: nucleus accumbens
AD: anterodorsal nucleus
AI: agranular insular area
AId: dorsal AI
AIv: ventral AI
AIp: posterior AI
AM: anteromedial nucleus
AOB: accessory olfactory bulb
AON: anterior olfactory nucleus
AUD: auditory areas
AV: anteroventral nucleus of the thalamus
BA: bed nucleus of the accessory olfactory tract
BAC: bed nucleus of the anterior commissure
BLA: basolateral amygdalar nucleus
BMA: basomedial amygdalar nucleus
BMAa: anterior BMA
BMAp: posterior BMA
BST: bed nuclei of the stria terminalis
CEA: central amygdalar nucleus
CEAc: capsular CEA
CEAl: lateral CEA
CEAm: medial CEA
CL: central lateral nucleus of the thalamus
CM: central medial nucleus of the thalamus
CLA: claustrum
COA: cortical amygdalar area
CP: caudoputamen
CTX: cerebral cortex
CTXsp: cortical subplate
DP: dorsal peduncular area
ECT: ectorhinal area
EP: endopiriform nucleus
FRP: frontal pole (cerebral cortex)
FS: fundus of striatum
GPe: globus pallidus, external segment
GPi: globus pallidus, internal segment
GRN: gigantocellular reticular nucleus
GU: gustatory areas
HIP: hippocampal region
HPF: hippocampal formation
HY: hypothalamus
IA: intercalated amygdalar nucleus
IAD: interanterodorsal nucleus of the thalamus
IAM: interanteromedial nucleus of the thalamus
IC: insular cortex
ILA: infralimbic area
ILM: intralaminar nuclei of the dorsal thalamus
IMD: intermediodorsal nucleus of the thalamus
IO: inferior olivary complex
IRN: intermediate reticular nucleus
LA: lateral amygdalar nucleus
LAT: lateral group of the dorsal thalamus;
LD: lateral dorsal nucleus of the thalamus
LH: lateral habenula
LHA: lateral hypothalamic area
LP: lateral posterior nucleus of the thalamus
LS: lateral septal nucleus
LSS: lateral strip of striatum
MA: magnocellular nucleus
MB: midbrain
MD: mediodorsal nucleus of thalamus
MDRN: medullary reticular nucleus
MEA: medial amygdalar nucleus
MED: medial group of the dorsal thalamus
MG: medial geniculate complex
MO: motor areas
MOp: primary motor area
MOs: secondary motor area
MOB: main olfactory bulb
MRN: midbrain reticular nucleus
MSC: medial septal complex
MTN: midline group of the dorsal thalamus
MY: medulla
NLOT: nucleus of the lateral olfactory tract
NTS: nucleus of the solitary tract
OLF: olfactory areas
ORB: orbital area
OT: olfactory tubercle
P: pons
PA: posterior amygdalar nucleus
PAA: piriform-amygdalar area
PAG: periaqueductal gray
PAL: pallidum
PARN: parvicellular reticular nucleus
PB: parabrachial nucleus
PCN: paracentral nucleus
PERI: perirhinal area
PF: parafascicular nucleus
PG: pontine gray
PIR: piriform area
PL: prelimbic area
PO: posterior complex of the thalamus
PR: perireunensis nucleus
PT: parataenial nucleus
PTLp: posterior parietal association areas
PVT: paraventricular nucleus of the thalamus
RE: nucleus of reuniens
RH: rhomboid nucleus
RHP: retrohippocampal region
RSP: retrosplenial area
RT: reticular nucleus of the thalamus
SCm: superior colliculus, motor related
SF: septofimbrial nucleus
SH: septohippocampal nucleus
SI: substantia innominata
SMT: submedial nucleus of the thalamus
SNc: substantia nigra, compact part
SNr: substantia nigra, reticular part
SS: somatosensory areas
SSp: primary SS
SSs: supplemental SS
STR: striatum
TEa: temporal association areas
TH: thalamus
TR: postpiriform transition area
TRN: tegmental reticular nucleus
TRS: triangular nucleus of septum
TT: taenia tecta
VAL: ventral anterior-lateral complex of the thalamus
VENT: ventral group of the dorsal thalamus
VIS: visual areas
VISC: visceral area
VM: ventral medial nucleus of the thalamus
VP: ventral posterior complex of the thalamus
VTA: ventral tegmental area
ZI: zona incerta

## Methods

### Mice

All experimental procedures were approved by the Animal Care and Use Committee of the Center for Excellence in Brain Science and Intelligence Technology (Institute of Neuroscience), Chinese Academy of Sciences. Male C57BL/6J mice (Shanghai Experimental Animal Center) were used for all experiments. Animals were housed under standard laboratory conditions with a 12-h light/dark cycle, ambient temperature maintained at 22–25 °C, and relative humidity between 30– 70%, with food and water provided ad libitum.

### Single-cell projectome data acquisition and analysis

Male C57BL/6J mice (8–10 weeks old) were group-housed prior to surgery. Mice were anesthetized with isoflurane, and adeno-associated viruses (AAVs) were injected into the insular cortex (IC), including the agranular insular area (AI), gustatory area (GU), and visceral area (VISC), to achieve sparse neuronal labeling. Whole-brain imaging was performed using fluorescence micro-optical sectioning tomography (fMOST) as previously described^60,61^. Each dataset contained two imaging channels: EGFP, used for neurite tracing, and propidium iodide (PI), used for anatomical registration. Neurons were reconstructed by integrating tracings from two independent annotators and verified by a third for quality control. A composite scoring system incorporating multiple error metrics was applied to ensure reconstruction accuracy. All validated neurons were spatially registered to the Allen Mouse Brain Common Coordinate Framework (CCFv3). Subsequent analyses were performed using the NeuronVis Python package and custom Python scripts for quantification, visualization, and network analysis.

### Clustering of single-neuron projectome

To classify IC neurons based on their whole-brain projection patterns, we applied the FNT-dist algorithm to quantify pairwise morphological dissimilarities among all reconstructed neurons^62^. The resulting dissimilarity matrix was subjected to hierarchical clustering using Ward’s linkage method to delineate distinct projectome-based neuronal subtypes. The optimal number of clusters was determined by evaluating the silhouette score (range: -1 to 1), computed with the silhouette_score() function in scikit-learn package, where higher values indicate greater clustering coherence. Twelve neuronal subtypes were selected corresponding to the local maximum of the silhouette score.

### Entropy of soma distribution

To quantify the spatial specificity of soma distribution for each neuronal cluster across IC subregions, we calculated an entropy index (*S*_*i*_) as follows:

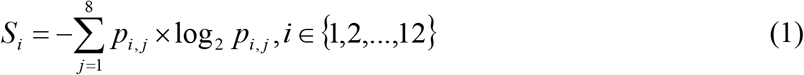

where *p*_*i,j*_ represents the proportion of neurons in cluster *i* whose somata are located within subregion *j* to the total number of neurons in that cluster. A higher *S*_*i*_ indicates a more dispersed spatial distribution across IC subregions, whereas a lower *S*_*i*_ reflects greater spatial concentration within a specific subregion.

### Contour map of neuron density

To correct for variability introduced by sparse labeling and non-uniform sampling across IC subregions, we normalized the spatial distribution of neuronal somata and visualized their density using contour maps. Soma coordinates were first projected onto a two-dimensional plane defined by the anteroposterior (AP) and mediolateral (ML) axes. The plane was then divided into grids, and for each grid, the proportion of neurons belonging to a given subtype was calculated as the probability of occurrence within that region. Assuming an equal total neuron count per grid (e.g., 1,000 neurons), corresponding soma coordinates were randomly generated according to these probabilities. Kernel density estimation (KDE) was then applied to the simulated coordinates using the kdeplot() function in the Seaborn package, producing a continuous contour representation of neuron density across the IC.

### Axon projection intensity analysis

Projection intensity within a target region was quantified as either the number of axonal terminals or the total axon length, both of which correlate with the density of synaptic innervation. To compare across regions, projection intensity was normalized by dividing the total number of terminal points contributed by all neurons within a given region of interest by the total number of terminal points observed across the entire brain. This normalized value represents the relative strength of axonal projection from IC neurons to each target region. For comparing projection intensities among multiple subregions within a brain region–such as the subnuclei of the amygdala–projection intensity was normalized as follows: the number of terminals from a population of IC neurons in a specific amygdala subregion was divided by the total number of terminals from the same population across the entire amygdala.

### Decomposing a neuron into primary axonal trunk, first-order axonal collaterals, and high-order branches

To analyze the hierarchical organization of axonal branching, each reconstructed neuron was decomposed into a primary axonal trunk, first-order collaterals, and higher-order branches based on geometric branching rules. The axonal path extending from the soma to its terminal endpoint was defined as the primary axonal trunk (order 0). Subsequent branch orders were determined according to the branching angles at each bifurcation. Branches were classified as symmetric when both daughter branches diverged at angles greater than 45° or when the ratio of their branching angles was less than 1.7; in such cases, both daughter branches were assigned an order one level higher than that of the parent branch. When these criteria were not met, the bifurcation was defined as collateral: the daughter branch with the smaller branching angle inherited the same order as the parent, while the other branch was assigned an order incremented by one.

### Clustering of first-order axonal collaterals

First-order axonal collaterals were defined as branches of order 1 whose corresponding tree compartments contained at least 150 tracing points. To classify these collaterals based on their projection morphology, we computed pairwise morphological distances using the FNT-dist algorithm, generating a dissimilarity matrix analogous to that used for whole-neuron clustering. Hierarchical clustering with Ward’s linkage method was applied to this matrix to identify distinct collateral groups. The silhouette score was calculated to evaluate cluster quality, and the number of clusters corresponding to the local maximum of the silhouette score was selected as the optimal cluster number.

### Quantification of collateral contribution to total projection intensity

To quantify the contribution of axonal collaterals to total projection intensity, and thereby evaluate the degree of expansion of neuronal innervation territory mediated by these collaterals, we defined an expansion rate metric and derived its mathematical formulation:

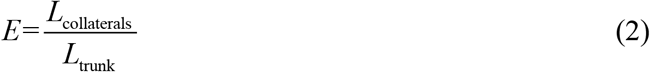

where *E, L*_collaterals_, and *L*_trunk_ represent expansion rate, cumulative length of all axonal collaterals, and length of primary axonal trunk, respectively.

### Probability density of neuronal somata along the AP axis

To visualize the probability density of soma distributions along the AP axis, each IC region was divided into bins with a step size of 0.25 mm. For each bin, the number of neurons belonging to a specific subtype was divided by the total number of neurons in that region to obtain the relative frequency, which was treated as the probability of occurrence within that bin. Assuming a uniform distribution of neurons across bins (1,000 neurons per bin), we multiplied this constant by the calculated probability to generate the normalized neuron count for each bin. Corresponding AP coordinates were then randomly generated within each bin according to these counts. KDE was applied to the simulated AP coordinates using the kdeplot function from the Python package Seaborn, producing a continuous probability density profile of soma distribution along the AP axis.

### Linear regression analysis

Linear regression analyses were performed using the regplot() function from the Seaborn package to fit and visualize regression lines. The correlation coefficient (*r*) and p-value were computed using the linregress() function from the SciPy library to assess the strength and significance of the linear relationship.

### Dendritic orientation relative to laminar direction

Let the spatial coordinates of the soma be A, and the spatial coordinates of any point on the dendrite be B. Thus, a three-dimensional vector 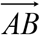 can be obtained. By summing the vectors formed by pairing coordinates of all sampling points on the dendrite with the soma coordinates, the sum vector of the dendrite (**v**_sum_) can be derived. The angle between the dendritic sum vector and the unit vector that is perpendicular to the cortical surface and originates from the soma (**v**_laminar_) represents the dendritic orientation relative to the laminar direction (*θ*), which can be calculated based on the dot product formula:

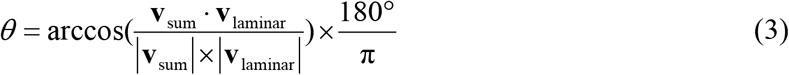

### Construction of IC units

A K-means clustering–based algorithm was applied to segment the surface of the IC into 40 spatially contiguous patches of approximately equal area. To extend these surface patches through the full cortical thickness, we adopted the streamline-based volumetric expansion method developed by the Allen Institute^63^, which propagates each surface patch along the laminar (surface-normal) direction to generate complete three-dimensional IC units for subsequent connectivity analyses.

### Identification of modular structures in the intra-IC network

Each reconstructed IC neuron was assigned to one of the predefined IC units based on its soma coordinates. The projection intensity from a source unit to a target unit was defined as the mean axonal length of all neurons originating in the source unit and terminating in the target unit, forming a weighted intra-IC connectivity matrix. To identify modular organization within this network, we applied the Louvain community detection algorithm implemented in the Python package bctpy. The resolution parameter (γ), which controls the granularity of module detection, was systematically varied from 0.05 to 0.9 in increments of 0.05. For each γ value, modularity optimization was repeated 1,000 times for both the original and a shuffled version of the intra-IC connection matrix. The optimal modular partition was determined at γ = 0.9, where the difference between the modularity values of the true matrix (*Q*_True_) and the shuffled matrix (*Q*_shuffle_) reached its maximum, indicating the most stable and biologically meaningful modular structure.

### Intra-IC network graph

To visualize intra-IC connectivity, network graphs were generated using the Python package NetworkX. The geometric center of each IC unit’s surface was used to represent the corresponding node. Directed edges were drawn between nodes when the projection intensity from a source unit to a target unit, as defined in the intra-IC connection matrix, was ≥ 0.4; weaker connections were omitted from visualization. This thresholded representation highlights the principal structural pathways within the intra-IC network.

### Convergence index

To quantify each IC unit’s tendency to integrate or disseminate information within the intra-IC network, we defined a convergence index (CI). For a given unit, let denote the number of other units projecting to it with projection intensity above a defined threshold, and *N*_output_ the number of units it projects to with intensity above the same threshold. The convergence index was computed as:

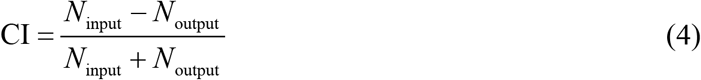

Because CI depends on the projection intensity threshold, values ranging from 0.21 to 0.81 (step size, 0.01) were tested. A CI was calculated for each threshold, and the resulting values were evaluated using a one-sample t-test. Units with CI values significantly greater than 0.5 were classified as convergent nodes, those significantly less than -0.5 as divergent nodes, and those within this range as recurrent nodes.

### Rate-based whole-IC model

We constructed a rate-based whole-IC network model in which each IC unit was represented by a FitzHugh–Nagumo (FHN) oscillator. Units were interconnected according to the empirically derived intra-IC connectivity matrix, with coupling strength proportional to projection intensity. Signal transmission delays between units were estimated from the mean axonal length of the corresponding projections and incorporated into the model dynamics. All IC units were coupled using a diffusive coupling framework, allowing the network to capture both local and long-range interactions. The simulated constant current inputs were applied to the network to mimic external sensory stimulation, following a temporal pattern of “1 s baseline, 1 s stimulus, and 2 s post-stimulus window.” Simulations were implemented in BrainPy^64^ and run for 4,000 ms with a 0.1-ms time step. The resulting time series of unit activity were analyzed by computing pairwise Pearson’s correlation coefficients to generate a functional connectivity matrix. The Louvain algorithm was then applied to this matrix to identify the optimal set of functional modules within the IC network.

### Construction and training of recurrent neural network (RNN)

Recurrent neural networks (RNNs) were implemented using BrainPy and PyTorch, with the latter combined with the loss-landscape package for analysis of the optimization surface. The hidden layer contained 40 recurrent units, corresponding to the 40 IC subdomains defined by the intra-IC connection matrix. Units were spatially allocated according to anatomical proportions—13 units to the aIC, 14 units to the pIC, and the remaining units to the mIC. The input dimension was set to 400, representing auditory cortical neurons projecting to the IC. Based on known anatomical pathways, pIC units served as network input nodes and aIC units as output nodes. To enforce biologically inspired connectivity, input and output masks were applied during training. The input mask restricted gradient updates to weights targeting pIC units, while the output mask, applied at initialization, limited dense-layer connectivity to aIC units. Only input weights, recurrent weights, and biases were trainable; output weights and biases remained fixed. All RNNs used rectified linear unit (ReLU) activation functions. Hidden states from pIC units were passed to a dense output layer producing a two-dimensional output at each timestep. Outputs across time were averaged by global temporal pooling, and the resulting vector was compared with ground-truth labels using cross-entropy loss. Classification accuracy was computed as the proportion of samples whose predicted class (maximum output index) matched the true label. Input spike data were generated using a custom Poisson population module in BrainPy, where each neuron’s activity at each 1-ms timestep was binary (1 = spike, 0 = no spike). For go and no-go trials, the mean firing rates of auditory cortical neurons were set to 3 Hz and 10 Hz, respectively, with variances of 0.002 Hz and 0.1745 Hz, consistent with a previous study^65^. Each neuron had a 50-ms refractory period following a spike. Each sample consisted of 400 neurons spiking over 300 ms, and the dataset included 2,048 training and 512 test samples. Training was performed using backpropagation through time (BPTT) for 100 epochs, with the AdamW optimizer (weight decay = 1 × 10^−4^) and a cosine-annealing learning-rate schedule, decaying from 1 × 10^−4^ to 1 × 10^−6^ over the course of training. Networks were trained in mini-batches of 64 samples, randomly drawn from the training dataset at each iteration.

### RNN initialization

To investigate how intra-IC connectivity patterns influence learning efficiency in RNNs, we implemented four distinct initialization schemes for the recurrent weight matrix (**W**) in the go/no-go task:

#### (1) Insula-origin initialization

The scaled intra-IC connection matrix (**W**_scaled_) was perturbed with controlled multiplicative Gaussian noise to simulate biologically plausible inter-individual variability while preserving the underlying topological organization of intra-IC connectivity:

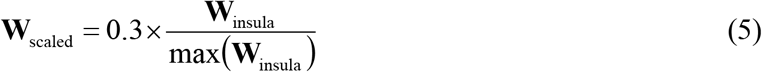

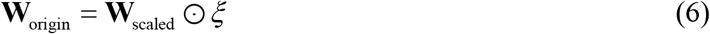

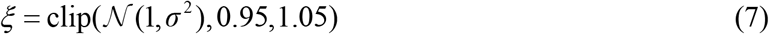

where **W**_insula_ represents the original intra-IC connection matrix, ⊙ denotes Hadamard (element-wise) product, and the random variable *ξ* follows a clipped Gaussian distribution with mean 1 and variance *σ*^2^ (*σ* = 0.02). This controlled perturbation preserved the overall topological structure of intra-IC connectivity while introducing realistic variability across simulated individuals.

#### (2) Insula-shuffle initialization

Elements of **W**_scaled_ were randomly permuted using the weighted degree-preserving shuffling method. Firstly, two elements at different positions in **W**_scaled_ were randomly selected, denoted as *W*_*i,l*_ and *W*_*k,j*_, representing the values in row *i* column *l* and row *k* column *j*, respectively. Since **W**_scaled_ contains no negative values, to prevent negative entries after shuffling, *δ*_upper_ was defined as the minimum of *W*_*i,l*_ and *W*_*k,j*_. A random value *δ* was then sampled from a uniform distribution over [0, *δ*_upper_]. Subsequently, the elements of **W**_scaled_ were updated according to the following equations:

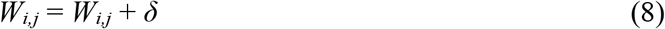

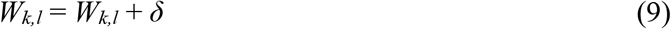

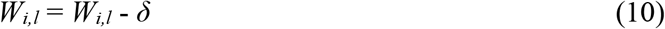

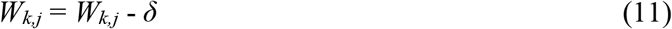

To ensure thorough shuffling of **W**_scaled_, the above procedure was repeated 100,000 times. This process disrupted higher-order topological structure of **W**_scaled_, including clustering, motif patterns, and path organization, while strictly preserving the weighted in- and out-degree of each node. The resulting **W**_shuffle_ thus served as a control to isolate the contribution of weighted degree sequence from other topological features in shaping RNN learning dynamics.

#### (3) Norm-preserved random initialization (Random-Fro)

Random matrices were generated using Kaiming-normal initialization. After taking absolute values (to ensure non-negativity, consistent with **W**_scaled_), the resulting matrices were rescaled to match the Frobenius norm of **W**_scaled_:

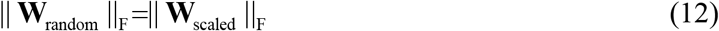

This ensured that all networks shared comparable overall weight magnitudes while differing in topological organization.

#### (4) Spectrum-preserved random initialization (Random-SR)

Random matrices were generated using Kaiming-normal initialization. After taking absolute values (to ensure non-negativity, consistent with **W**_scaled_), the resulting matrices were rescaled to match the spectral radius of **W**_scaled_:

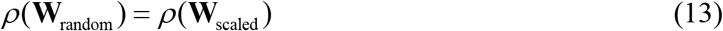

This preserved the global stability and dynamic range of the recurrent network while removing the specific structural correlations of the IC topology.

For each random seed, all four initialization schemes were applied to generate 20 independent recurrent weight matrices per condition. At each seed, input and output weight matrices were initialized independently using Kaiming-normal distribution, and all bias terms in both the RNN and output layer were set to zero. To ensure biological plausibility, all input and output weights were transformed to their absolute values. Finally, to reflect the terminal distribution of auditory cortical inputs within the IC, input weights corresponding to specific pIC units (p2, p9, p20, p35) were selectively enhanced, whereas those of pIC units (p8, p12, p17, p19, p26, p34, p39) were attenuated during initialization.

### Somatosensory cortical network

To investigate whether the internal connectivity pattern of other brain regions beyond the IC could similarly accelerate RNN learning, we selected the somatosensory cortex (SS) as a test case. The SS was partitioned into 40 units, based on which its internal connection matrix was derived. The recurrent weights of the SS-based RNNs were then initialized using the same methodology applied to the intra-IC connection matrix:

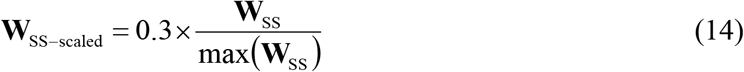

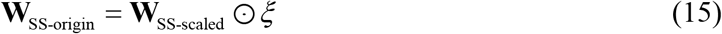

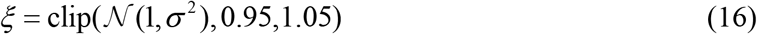

where *W*_*SS*_ represents the original intra-SS connection matrix, ⊙ denotes Hadamard (element-wise) product, and the random variable *ξ* follows a clipped Gaussian distribution with mean 1 and variance *σ*^2^ (*σ* = 0.02). For the SS-based RNNs, input and output nodes were assigned following a typical biological pathway (SSp→SSs): nodes corresponding to SSp served as the input nodes for external information, while nodes corresponding to SSs were designated as the output nodes of the RNNs.

### Generalization of IC architecture across diverse tasks

To evaluate the IC architecture in a broader range of contexts, we selected five representative classic tasks for testing. To achieve a general RNN training conditions, we removed constraints on input and output nodes and allowed the output layer to participate in gradient updates. Furthermore, minor adjustments were made to the initialization of recurrent weights: for the Insula-origin and Random-SR groups, weight matrices were scaled to a target spectral radius of 0.8; for the Random-Fro and Random-SR groups, weight matrices were generated using Xavier-normal initialization (to match the tanh activation function); all other initialization steps remained consistent with those used in the go/no-go task. Besides, the training protocol remained identical to that of the go/no-go task: only the initialization of the recurrent weights differed between RNN groups, while all other initial conditions and hyperparameter settings were kept consistent.

#### (1) Delayed match-to-sample task

In this task, the RNN agent was required to memorize the first sample and, after a delay, compare it with a second sample to determine if they matched. Each trial’s input was a 12×10 matrix, where each column represented an input channel and each row a timestep. The first and second samples appeared at the first and twelfth timesteps, respectively. To construct the input, the first sample was randomly drawn from a standard normal distribution. In match trials, the second sample was identical to the first; in non-match trials, the second sample was independently drawn from a standard normal distribution. Finally, Gaussian noise drawn from a normal distribution with mean μ = 0 and standard deviation σ = 0.1 was added to the entire input matrix. In match trials, the target was set to 1, while in non-match trials it was set to 0. The training set consisted of 2048 trials, and the validation set contained 512 trials. The RNN input, hidden, and output dimensions were set to 10, 40, and 1, respectively. The activation function was tanh. The network output at the final timestep was passed through a linear readout layer, followed by a sigmoid function to produce the final output. The model was trained using binary cross-entropy (BCE) loss and optimized with AdamW (weight decay = 1 × 10^−4^). A cosine annealing schedule was employed, with an initial learning rate of 1 × 10^−3^ and a minimum learning rate of 5 × 10^−4^. Gradient clipping was applied to all parameters with a maximum norm of 1.0. A batch size of 64 was used, and training proceeded for 100 epochs. Experiments were run across 10 different random seeds, with 20 independent runs per seed, yielding a total of 200 training runs for statistical analysis.

#### (2) Interval reproduction task

The task required the RNN agent to reproduce the time interval between cue 1 and cue 2 by triggering a Go response after cue 2, aiming to match *t*_Go_ -*t*_cue2_ to *t*_cue2_ -*t*_cue1_. On each trial, the network received two discrete input pulses, cue 1 at *t*_cue1_ = 0 and cue 2 at *t*_cue2_, which were separated by a variable interval drawn uniformly from the integers 20 to 60. The network needed to make the Go response at *t*_Go_ = 2 × *t*_cue2_ by generating a Gaussian target output centered at *t*_Go_ with standard deviation *σ* = 5.0. A binary mask excluded loss computation before the cue 2 (*t* ≤ *t*_cue2_) to prevent anticipatory responses. Each sequence comprised 150 time steps with input and target dimensions of 1. The training set consisted of 2048 trials, and the validation set contained 512 trials. The RNN had input, hidden, and output dimensions of 1, 40, and 1, respectively, and used tanh as the activation function. The network outputs from all timesteps were linearly read out to produce the final output. The model was trained with mean squared error (MSE) loss, optimized using AdamW (weight decay = 1 × 10^−4^), and scheduled with cosine annealing (initial learning rate = 1 × 10^−3^, minimum learning rate = 1 × 10^−5^). Gradient clipping was applied to all parameters with a maximum norm of 1.0. Training was conducted for 100 epochs across 10 different random seeds, with 20 independent runs per seed, yielding a total of 200 runs for statistical analysis.

#### (3) Context-dependent task

In this task, the RNN agent was required to adapt its decision policy according to the context. Each trial comprised three periods: context (10 steps), stimulus (10 steps), and response (5 steps). Context was signaled by activation of one of two dedicated input channels (context A or B), followed by simultaneous presentation of two noisy stimulus values drawn from an anti-correlated distribution (*s*_B_ = −*s*_A_ + 𝒩 (0, 0.05^2^)). The RNN agent was required to report which of the two stimuli had the larger signed value, but only for the stimulus dimension specified by the context cue. Loss was computed exclusively during the response period. The training set consisted of 4096 trials, and the validation set contained 1024 trials. The RNN had input, hidden, and output dimensions of 4, 40, and 2, respectively, with tanh as the activation function. A linear readout was applied to the outputs of all timesteps to produce the final prediction. Training used cross-entropy loss and the AdamW optimizer with weight decay of 1 × 10^−4^, under a constant learning rate of 3 × 10^−4^. Gradient clipping was applied to all parameters with a maximum norm of 1.0. Training proceeded for 100 epochs across 10 different random seeds, with 20 independent runs per seed, resulting in a total of 200 runs for statistical analysis.

#### (4) Four-armed bandit task

In this classic reinforcement-learning task, the RNN agent was required to explore four slot machines with fixed reward probabilities of 0.6, 0.55, 0.5, and 0.45. On each trial, the agent selected one of the four machines and received a reward (coded as 1) or no reward (coded as 0). The agent’s goal across 3000 trials was to maximize cumulative reward. The input on each trial was a 5-dimensional vector: the first four dimensions indicated the agent’s action on the previous trial (e.g., (0, 1, 0, 0) corresponded to selecting the second machine), and the fifth dimension indicated whether a reward was received (1 for reward, 0 for no reward). The RNN had input, hidden, and output dimensions of 5, 40, and 4, respectively, using tanh activation and a linear readout to produce the final output. Training employed the AdamW optimizer with weight decay of 1 × 10^−4^ and the REINFORCE algorithm for policy loss. Action sampling was optimized using Gumbel-Softmax with a temperature that decayed exponentially from 5.0 to 0.3. A constant learning rate of 3 × 10^−4^ was used, and gradient clipping was applied to all parameters with a maximum norm of 1.0. The model was trained for 100 epochs across 10 random seeds, with 20 independent runs per seed, yielding 200 runs in total for statistical analysis. The reward was smoothed using an exponentially weighted moving average, computed as *reward* = 0.05 × *r*_*t*_ + 0.95 × *reward*, where *r*_*t*_ denotes the immediate reward at current trial *t*. Regret was defined as the difference between the reward probability of the chosen action and the maximum reward probability available in a trial. The optimal action rate was calculated as the proportion of trials, within a sliding window of the most recent 100 trials, in which the agent selected the option with the highest reward probability. The AUC for reward, regret, and optimal action rate was computed on raw data, whereas the corresponding curves were smoothed using a Savitzky-Golay filter (window length = 51, polynomial order = 1).

#### (5) Penn Treebank (PTB) language modeling

This task serves as a benchmark for assessing an RNN’s ability to capture long-range dependencies and sequential regularities. The RNN agent must learn contextual-historical dependency patterns to acquire syntactic and semantic knowledge, and then predict the probability distribution of the next word given the preceding word sequence. To construct the training dataset, sentences were extracted from the PTB using the Natural Language Toolkit (NLTK). Input sequences were constructed by tokenizing the corpus into words, lowercasing, and mapping to a vocabulary of the 10,000 most frequent tokens (plus <pad> and <unk> special tokens). The dataset was partitioned into non-overlapping sequences of 35 tokens, with each input sequence paired with its right-shifted counterpart as the prediction target. Models were trained with a batch size of 64 using contiguous sequential batches without shuffling. The RNN was configured with input, embedding, hidden, and output dimensions of 10,000, 128, 40, and 10,000, respectively. The embedding layer was kept frozen across all training runs and did not receive gradient updates. The hidden layer used a tanh activation, and the output layer was linear. The model was trained using the AdamW optimizer with a weight decay of 1 × 10^−4^, cross-entropy loss, a fixed learning rate of 3 × 10^−3^, and gradient clipping applied to all parameters with a maximum norm of 5.0. Training was conducted over 10 random seeds, with 20 independent runs per seed, each run lasting 100 epochs, yielding a total of 200 training runs for statistical analysis.

Perplexity (PPL) was computed as the exponential of the average negative log-likelihood (cross-entropy loss) over all tokens:

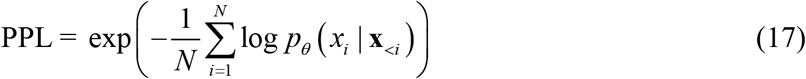

where *p*_*θ*_ denotes the model’s predicted probability and *N* is the total number of tokens in the test dataset.

Top-5 accuracy was defined as the proportion of test tokens for which the ground-truth word appears among the model’s five highest-probability predictions:

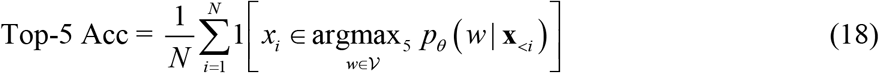

where *w* and *v* represents token and vocabulary, respectively. 1[·] is the indicator function.

### Random seeds and reproducibility

To minimize stochastic variability, 10 independent random seeds (42, 860, 3207, 8672, 15795, 37194, 44131, 56472, 60820, 87289) were used for training runs in all tasks. An additional set of random seeds (0-9) was used to control environmental interactions in the four-armed bandit task, with each environmental seed paired one-to-one with the corresponding training seed. Before each RNN training batch, the computational environment was deterministically initialized with the designated seed to ensure full reproducibility of all random operations, including weight initialization and data sampling. Results from all seeded trials were subsequently aggregated for statistical analyses, thereby controlling for seed-dependent fluctuations and enabling ensemble-level evaluation of model performance.

### Recurrent weights trajectory and clusterability

During training, all recurrent weight matrices of the RNN were recorded, flattened into vectors, and concatenated to form a two-dimensional matrix representing weight evolution across epochs. t-SNE dimensionality reduction was then applied to visualize the temporal trajectory of recurrent weights throughout training. For each initialization method and random seed, 20 independent training runs were conducted, each producing a distinct trajectory in the reduced-dimensional space. To quantify trajectory similarity, pairwise Fréchet distances were computed among the 20 trajectories, generating a distance matrix that served as input for hierarchical clustering using Ward’s linkage method. Clustering was evaluated over a range of candidate cluster numbers (up to 10), and the silhouette score was computed for each partition. The maximum silhouette score obtained across this range was taken as a quantitative measure of trajectory clusterability for the corresponding initialization condition.

### Loss surface and Hessian matrix

The loss-landscape package^66^ and software Paraview were employed to compute and visualize the loss surface, allowing analysis of the RNN’s convergence properties by characterizing the distribution of loss values across parameter space. The Hessian matrix (**H**), representing the second-order derivatives of the loss function with respect to model parameters, captures the local curvature of the optimization landscape through its eigenvalue spectrum. The spectral radius and condition number of the Hessian were defined as:

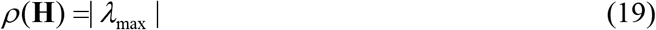

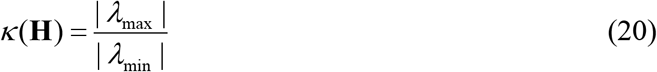

where *ρ*(**H**), *κ*(**H**), *λ*_*max*_, and *λ*_*min*_ represent the spectral radius, condition number, maximum and minimum eigenvalues of the Hessian matrix, respectively. *ρ*(**H**) characterizes the curvature of the loss function along its steepest direction at the current parameter point, with larger values indicating more pronounced curvature fluctuations and consequently reduced optimization stability. *κ*(**H**) describes the degree of variation in curvature across different parameter directions, thereby quantifying the difficulty of the optimization problem. A larger *κ*(**H**) indicates greater disparity in curvature across different directions (higher anisotropy), potentially leading to slower convergence of optimization algorithms.

### Low-rank tensor decomposition

To uncover latent patterns in recurrent weight dynamics during RNN training, all recurrent weight matrices—across initialization methods, random seeds, repeated runs, and training epochs—were flattened into vectors and assembled into a fifth-order tensor χ ∈ ℝ^*M*× *S*×*R*×*E*×*W*^, where *M, S, R, E*, and *W* respectively represent method, seed, run, epoch, and flattened weight. This high-dimensional tensor was subsequently factorized using a R-component CP (CANDECOMP/PARAFAC) decomposition model, expressed as:

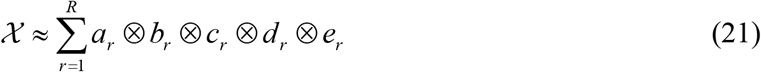

where ⊗ represents the vector outer product, *a*_*r*_ ∈ ℝ^*M*^, *b*_*r*_ ∈ ℝ^*S*^, *c*_*r*_ ∈ ℝ^*R*^, *d*_*r*_ ∈ ℝ^*E*^ and *e*_*r*_ ∈ ℝ^*W*^, and *a*_*r*_ ⊗ *b*_*r*_ ⊗ *c*_*r*_ ⊗ *d*_*r*_ ⊗ *e*_*r*_ is a rank-one tensor. Before low-rank tensor decomposition using CP, *X* was normalized using z-score along the epochs and weights dimensions to improve the numerical stability, decomposition efficiency, and interpretability. CP decomposition was performed using the parafac function from the Python package TensorLy, employing the alternating least-squares (ALS) optimization algorithm.

To determine the optimal tensor rank for fitting CP models, we referred to the method described by Drieu et al^65^. In brief, for each candidate rank *R* ∈{1,…,6}, we performed *N* = 20 independent CP decompositions with random initializations (maximum 500 iterations, tolerance 10^−6^, normalized factors). The decomposition with minimum reconstruction error was designated as the reference model. Solution stability was quantified by computing pairwise component similarities between the reference model and all other runs. Due to the permutation indeterminacy of CP decomposition, components were aligned using a greedy matching algorithm based on cosine similarity across all modes. The mean similarity score across *N* - 1 comparisons served as the stability metric for rank *R*. In our data, a lower reconstruction error indicates better CP model fit, while a higher mean similarity score indicates greater stability. By jointly considering these two metrics, we ultimately selected *R* = 4 (Extended Data Fig. 8g).

The output of the decomposition was a set of four components, each composed of five factors: (1) method factor (*a*_*r*_), indicating static biases induced by initialization methods; (2) seed factor (*b*_*r*_), characterizing stochastic variations across different random number generator environments; (3) run factor (*c*_*r*_), quantifying reproducibility effects through repeated training instances; (4) epoch factor (*d*_*r*_), reflecting temporal evolution patterns during optimization; (5) weight factor (*e*_*r*_), embodying the intrinsic update strategies governing parameter adjustments.

### Average update magnitude (AUM)

To quantify the pace of recurrent-weight adaptation during the early training, we computed the L2-norm of weight changes after each epoch and averaged these values across the first *E* epochs (*E* = 18 in this study):

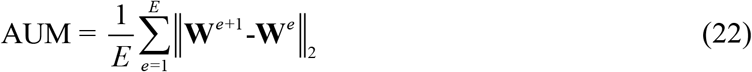

where *e* and **W** respectively denotes epoch *e* and the recurrent weight matrix, and ‖ ‖_2_ represents L2 norm. Higher AUM indicates more vigorous parameter exploration.

### Morphological characteristics of axons

An axonal branch point was defined as the location where an axon bifurcates into two or more daughter branches. The total axon length was calculated as the sum of the lengths of all branches within a neuron. For each branch, tortuosity was defined as the ratio between the arc length of the branch and the Euclidean distance between its start and end points. The mean tortuosity of a neuron was computed as the arithmetic average of tortuosity across all branches. A branch was classified as asymmetric if either (1) the divergence angle between a daughter branch and its parent branch was < 45°, or (2) the ratio between the maximum and minimum deviation angles of the daughter branches relative to the parent exceeded 1.7. The asymmetry ratio for each neuron was then calculated as the proportion of asymmetric branches relative to the total number of branches. Following axonal tree decomposition, each branch was assigned an order value, and the maximum branch order of a neuron was used as a measure of its axonal morphological complexity. To quantify the spatial coverage of axonal projections within a given target region, we used the volume fraction metric, defined as the ratio of the convex hull volume to the bounding box volume of the axonal point cloud, where higher values indicate broader spatial coverage.

### Ipsi-preference index

To quantify the projection bias of neurons with bilateral targets, we calculated the ipsi-preference index (*I*_ipsi_) based on the relative number of axonal terminals in the ipsilateral and contralateral target regions. For each neuron, let *N*_ipsi_ and *N*_contra_ denote the numbers of terminals in the ipsilateral and contralateral regions, respectively. The index was defined as:

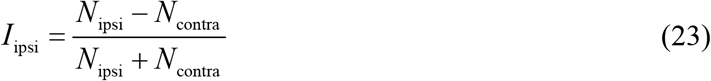

A significantly positive *I*_ipsi_ indicates a preference for ipsilateral projections, whereas a significantly negative value reflects a contralateral projection bias.

To assess each neuron’s overall hemispheric projection preference, a similar index was computed using the total terminal counts in the ipsilateral 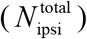 and contralateral 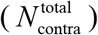 hemispheres:

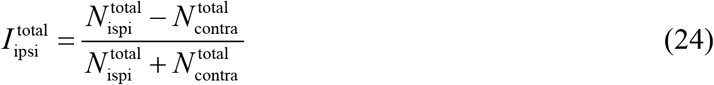

This global measure summarizes each neuron’s aggregate projection asymmetry across all target regions.

### Distribution of terminals along the AP axis

To quantify the spatial distribution of axonal terminals along the AP axis within a given brain region, terminal coordinates were binned at 0.1-mm intervals. The total number of terminals within each bin was counted, and the resulting histogram was smoothed using one-dimensional linear interpolation followed by a Savitzky–Golay filter to obtain a continuous distribution profile.

### Construction of the directed graph convolutional network (DGCN)

A directed graph convolutional network (DGCN) was implemented using the PyTorch Geometric (PyG) framework to extract topological features from the intra-IC connectivity matrices. The network architecture consisted of two graph convolutional layers. Input features were first processed through the initial convolutional layer, followed by a ReLU activation, and then passed to the second convolutional layer. The first and second layers produced 64-dimensional and 32-dimensional feature representations, respectively. Finally, a global mean pooling operation was applied to the output of the second layer to generate a compact 32-dimensional DGCN embedding representing the global topological structure of each connectivity matrix.

### Logistic regression classifier and confusion matrix

The DGCN embeddings obtained from the four matrix types (Insula-origin, Insula-shuffle, Random-Fro, and Random-SR) were pooled and randomly divided into training and test sets in a 4:1 ratio. A logistic regression classifier implemented in scikit-learn was trained to distinguish between the four matrix types based on their DGCN embeddings. Classification performance was evaluated using a confusion matrix, constructed by comparing the predicted and true labels on the held-out test set to assess overall prediction accuracy and misclassification patterns.

### Network analyses of connection matrices

Connection matrices were converted into directed graph representations using NetworkX^67^, and classical graph-theoretical analyses were performed. For these analyses, only the top 20% of strongest connections were retained to focus on the principal structural relationships, as existing evidence suggests that functionally predictive connections in the brain are strong connections, whereas weak connections contribute minimally to functional interpretation^68^. Weighted in-degree and out-degree metrics quantified the input and output capacities of each node, respectively. The average clustering coefficient was computed to assess the prevalence of local triangular motifs, reflecting the degree of interconnectivity among neighboring nodes. The average shortest path length served as an indicator of the cost of information transmission across the network. Global efficiency and local efficiency were calculated to evaluate, respectively, the overall robustness of information flow and the residual communication capacity among neighboring nodes after removal of individual nodes. The assortativity coefficient was used to examine connection similarity, indicating whether high-degree nodes preferentially connect to other high-degree nodes. Because the computation was performed on weighted directed adjacency matrices, we first removed scale differences using z^−^scoring, then mapped the values to the (0, 1) interval via a sigmoid function to ensure that differences in results were attributable to network topology rather than scale. To comply with the computational framework of NetworkX, we then took the reciprocal of the normalized adjacency matrix, converting it from a “projection intensity matrix” into a “cost/distance matrix.” All metrics were computed using methods adapted for directed graphs.

For spectral graph analyses, the complete (unthresholded) connection matrices were used. The normalized Laplacian spectra were computed, and KDE was applied to derive the spectral density distribution. The spectral gap (the second smallest eigenvalue of the Laplacian) was used as a measure of network modular separation. Additionally, adjacency matrix spectra were analyzed, and the first- and second-order spectral moment distances were calculated to quantify global coupling strength within the network. For motif analysis, again only the top 20% strongest connections were retained. Templates representing six common 3-node motifs and six common 4-node motifs were constructed. All possible 3-node and 4-node subgraphs were exhaustively enumerated, and only those exhibiting isomorphism with the predefined templates were included in motif counts.

### Identification of the most enriched motifs in the IC and SS

To identify the 10 most enriched motifs (considering only three- and four-node motifs) in the IC and SS, we employed the Python package igraph to search and identify motifs within the adjacency matrices. The top 10 motifs by occurrence count were selected as the final result.

### Identification of hubs within the IC

To identify key hub regions within the IC network, the intra-IC connection matrix was converted into a directed graph, and the PageRank algorithm was applied to quantify the relative importance of each IC unit in information propagation. The top 10 units with the highest PageRank scores were designated as IC hubs, representing nodes with dominant influence in intra-IC communication.

### Construction of networks with directed motif embeddings

To embed the specified motifs directionally into the network, we first initialized an *N* × *N* adjacency matrix with all elements set to zero, where N denotes the number of nodes in the network (*N* = 20 in this study). We then designated the first *n*_1_ nodes as input nodes and the last *n*_2_ nodes as output nodes (with *n*_1_ = *n*_2_ = 5 in this study). Next, nodes were randomly selected to form the specified motif, which was directionally embedded along the input–output axis into the adjacency matrix. For example, to embed a 4-node multi-layer feedforward motif, four nodes *a, b, c*, and *d* were randomly chosen and sorted in ascending order (assuming *a* < *b* < *c* < *d*). The elements at positions [*a,b*], [*a,c*], [*a,d*], [*b,c*], [*b,d*], and [*c,d*] in the adjacency matrix were then set to 1, completing one embedding operation. This process was repeated iteratively until either the target number of embeddings was achieved or the maximum allowable number of embedding attempts was reached. Embedding was skipped if the motif to be embedded was already present in the network. As a control for networks with directionally embedded motifs, we first counted the number of elements with a value of 1 (denoted as *m*) in the motif-embedded adjacency matrix. Subsequently, in a separate *N* × *N* adjacency matrix initialized with all zeros, *m* elements were randomly selected and set to 1 to construct the control network. Following this step, all elements with a value of 1 in both the motif-embedded and control adjacency matrices were replaced with random values drawn from a normal distribution with a mean of 1 and a standard deviation of 0.3, subject to a minimum value of 0.5. Then, positions with a value of 0 in these matrices were assigned random values sampled from a uniform distribution between 0.001 and 0.1, simulating weak connections in the brain. Finally, the resulting adjacency matrices were scaled according to their spectral radius to achieve a final spectral radius of 0.8.

### Comparison of the motif enrichment pattern

To compare the similarity of motif enrichment patterns among strong connections between the given two network types, we first quantified, for each network and its corresponding control within each type, the number of motifs formed by edges with connection strengths above the top 20%. The differences in these motif counts were then computed pairwise, resulting in a difference matrix. Subsequently, the mean along the observation dimension was calculated for the difference matrices of the two network types, yielding two vectors. A permutation test was employed to determine whether the observed Pearson correlation coefficient between these two vectors was significantly greater than that expected by chance, thereby assessing whether the motif enrichment patterns of strong connections were sufficiently similar across the two network types.

To further compare the similarity of motif enrichment patterns, we defined a pattern similarity score, as shown in the following formula:

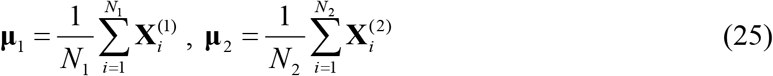

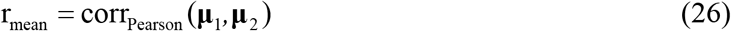

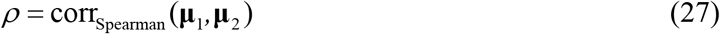

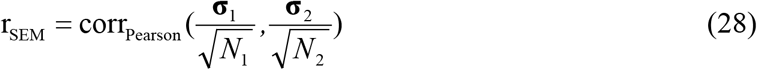

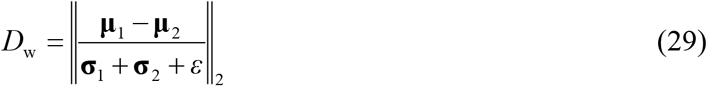

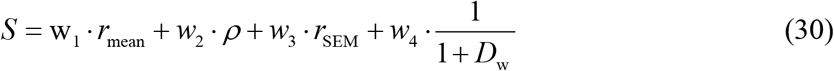

where 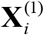 and 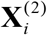 represent the difference matrices of two types of networks, *N*_1_ and *N*_2_ represent the sample sizes (number of observations) of 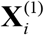 and 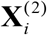, **μ**_1_ and **μ**_2_ are the mean vectors along the observation dimension, **σ**_1_ and **σ** _2_ are the standard deviation vectors along the observation dimension, *ε* = 10^*-*8^ is a small constant added for numerical stability, ‖ ‖_2_ represents L_2_ norm, and *w*_1_, *w*_2_, *w*_3_, and *w*_4_ are the weights used for weighted summation, with values of 0.3, 0.2, 0.1, and 0.4, respectively.

### Evaluation of network robustness

To assess network robustness, the adjacency matrix was incorporated as the recurrent weights of the RNN, which was trained to perform a go/no-go task. After successful training (i.e., when the RNN achieved a test accuracy above 0.95), the trained recurrent weights were extracted and subjected to perturbation. Two types of perturbations were applied: one involved randomly setting certain weights to zero (equivalent to deleting specific edges in the network), and the other involved randomly zeroing out all weights in a particular row and a particular column (equivalent to removing specific nodes). The perturbed recurrent weights were then used to replace the original trained weights in the RNN, and the network’s performance was re-evaluated on the test set to measure the decrease in accuracy.

In this study, both the experimental and control groups comprised 30 distinct RNNs, each initialized with a different recurrent weight matrix (derived from the network adjacency matrix). Each RNN was trained across 10 different random seeds. The robustness evaluation for each RNN was further conducted under 5 additional random seeds. The results were aggregated across these replications to mitigate the influence of stochasticity.

### Subdivision of the striatum based on IC inputs

To parcellate the striatum according to IC projections, the striatal volume, including both the caudoputamen and nucleus accumbens, was divided into 2,722 cubic voxels (200 × 200 × 200 μm^3^ each). For every striatum-projecting IC neuron, the axonal length within each voxel was quantified to construct a projection matrix representing voxel-wise innervation strength. Because some voxels contained no IC axons, minimal random perturbations were added to the projection matrix to eliminate zero values and allow computation of Spearman’s rank correlation coefficients between voxel pairs. Unsupervised hierarchical clustering was then performed using Spearman’s rank correlation as the distance metric and Ward’s linkage method, grouping voxels that received similar IC inputs. For each resulting cluster, the largest connected component was identified and smoothed using a Gaussian filter to generate the final striatal subregion. During integration of subregions, overlapping voxels were assigned based on penetration depth (distance to background), and residual gaps were filled by the nearest neighboring subregion. Each IC neuron was assigned to a striatal subregion if it exhibited the greatest cumulative axonal length within that subregion. The reconstructed striatal parcellation was visualized in three dimensions using the Mayavi package in Python.

### Terminal distribution map of neurons

To visualize the spatial distribution of axonal terminals within a specific coronal plane of a target region (e.g., the nucleus accumbens, ACB), terminal coordinates located within ±0.1 mm of the selected plane were extracted and projected onto that coronal plane for visualization. The anatomical boundary of the ACB at the corresponding level was delineated using binary erosion and overlaid onto the terminal map to provide anatomical reference.

### Hierarchy of the IC-thalamus network

The algorithms for computing hierarchical scores of IC units and thalamic subregions were adapted from established methods^62,69^. The initial hierarchical score of an IC unit or thalamic subregion *i* is mathematically expressed as:

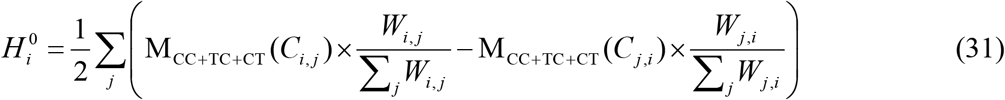

where *C*_*i,j*_ represents the projection pattern from unit j to unit i and M_CC+TC+CT_ serves as a composite mapping function defined based on a previous study^69^. For cortical-cortical (CC) projections, M_CC+TC+CT_ maps *C*_*i, j*_ to FF(=1) when *C*_*i, j*_ ∈{3,5,6} or FB(=-1) when *C*_*i, j*_ ∈{1,2,4}. Regarding thalamocortical (TC) projections, M_CC+TC+CT_ maps *C*_*i, j*_ to FF(=1) when *C*_*i, j*_ ∈{1,2,3,4} or FB(=-1) when *C*_*i, j*_ ∈{5}. For corticothalamic (CT) projections, linear discriminant analysis (LDA) was employed to classify them into either layer 5 (L5) or layer 6 (L6) dominance, thereby distinguishing FF from FB projections. The LDA was performed on log_10_-transformed number of IC units’ L5 and L6 neurons projecting to the thalamic subregions (including VENT, LAT, MED, MTN, and ILM), with the decision boundary constrained to approximate the y=x line. The *C*_*i, j*_ = 1 is assigned to projections with more L5 neurons, and *C*_*i, j*_ = 2 is assigned to L6 dominant projections. M_CC+TC+CT_ maps *C*_*i,j*_ to FF(=1) when *C*_*i, j*_ ∈{1} or FB(=-1) when *C*_*i, j*_ ∈{2}. It should be noticed that *C*_*i, j*_ = 0 when the projection intensity from unit *j* to unit *i* equals zero and M_CC+TC+CT_ consistently yields an output of 0 for all projection types (CC, TC, and CT) when *C*_*i, j*_ ∈{0}. *W*_*i, j*_ represents the mean projection intensity from unit *j* to unit *i*, operationally defined as the ratio of the total axon length in unit *i* originating from unit *j* to the total neuron number of unit *j*.

The final hierarchical score is iteratively computed using the following two equations until either the maximum iteration count (20) is reached or the norm of the difference between the current and previous final hierarchical score falls below the tolerance threshold (1×10^−6^):

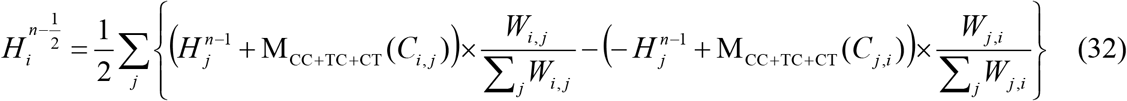

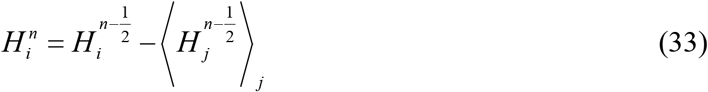

where *n* represents the number of iterative steps.

To evaluate the self-consistency of a hierarchy, the global hierarchy score is calculated as:

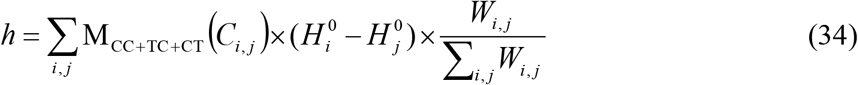

To assess statistical significance, shuffled connectivity matrices were generated, and the observed global hierarchy score was compared to the distribution of shuffled scores following the procedure described by the previous study^69^.

### Quantification and statistical analysis

All statistical analyses are explicitly stated where applicable. Statistical analysis was performed using GraphPad Prism 9 (GraphPad Software) and the Python package SciPy. The specific statistical tests employed for each comparison are detailed accordingly. Parametric tests were applied whenever the data met the assumptions of normality to compare means between two or more groups. For datasets with non-normal distributions, non-parametric tests were utilized. All statistical comparisons were conducted using two-tailed tests, with hypothesis testing performed at a significance threshold of 0.05 (n.s., *P* > 0.05; \**P* < 0.05; \*\**P* < 0.01; \*\*\**P* < 0.001; \*\*\*\**P* < 0.0001).

## Acknowledgements

We thank M.M. Poo and Y. Hu for reading the manuscript and helpful discussions; T.R. Xie for advice on calculating the thickness of each insular cortical layer; L. Gao for advice on FNT-dist implementation. This work was supported by grants from the Brain Science and Brain-like Intelligence Technology-National Science and Technology Major Project (2025ZD0215100 to X.X., 2025ZD0217800 to H.D.), CAS Project for Young Scientists in Basic Research (YSBR-114 to X.X.), National Natural Science Foundation of China (32371060 to X.X., 32271065, 32571197 to H.D.), Strategic Priority Research Program of the Chinese Academy of Sciences (XDB1010102 to X.X.), Lingang Laboratory (LG-QS-202203-02 to H.D.; LG-QS-202203-06 to X.X.), and Benyuan Charity Foundation (to H.D.).

## Author contributions

S.X., H.D., and X.X. conceived and designed the study. S.X. performed all data analyses. T.W., R.Z., X.Y.W., and R.S. contributed to data preparation and scientific discussion. X.F.W. provided guidance on projectome-related analyses. Y.C. and T.Z. advised on computational modeling. H.E. contributed to data interpretation and provided critical review and editorial input on the manuscript. S.X. and X.X. wrote the manuscript with contributions from all authors.

## Declaration of interests

The authors declare no competing interests.

## Figures, Extended Data, and Legend

**Extended Data Fig. 1.**
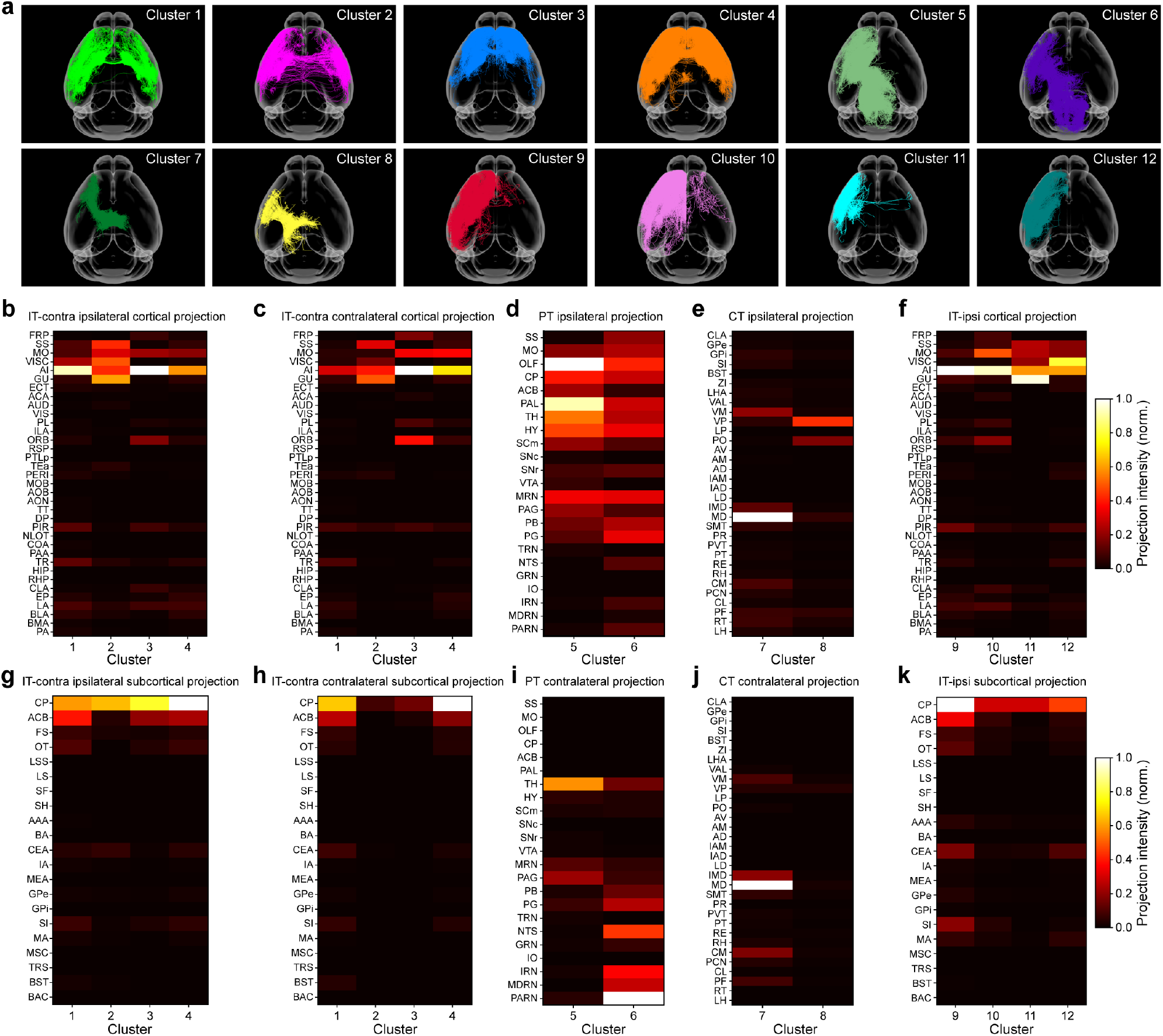
Classification and overall projection patterns of IC neurons. **a**, Whole-brain projectomes of neurons in each of the 12 clusters. **b**,**c** Normalized projection intensities of IT-contralateral clusters to ipsilateral (**b**) and contralateral (**c**) cortical targets. **d**, Normalized projection intensities of PT clusters to ipsilateral cortical targets. **e**, Normalized projection intensities of CT clusters to ipsilateral cortical targets. **f**, Normalized projection intensities of IT-ipsilateral clusters to ipsilateral cortical targets. **g**,**h** Normalized projection intensities of IT-contralateral clusters to ipsilateral (**g**) and contralateral (**h**) subcortical targets. **i**, Normalized projection intensities of PT clusters to contralateral targets. **j**, Normalized projection intensities of CT clusters to contralateral targets. **k**, Normalized projection intensities of IT-ipsilateral clusters to ipsilateral subcortical targets.

**Extended Data Fig. 2.**
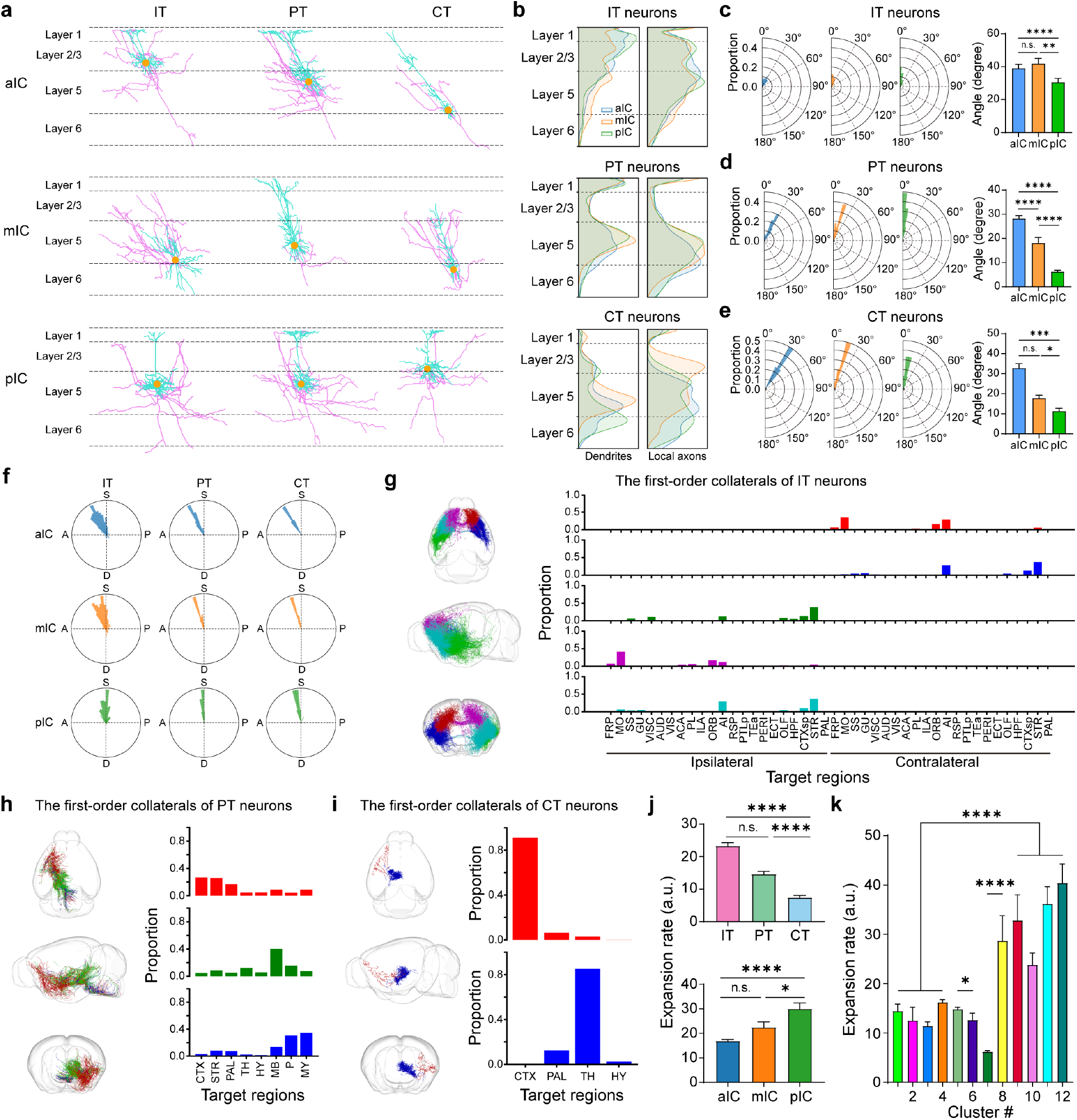
Morphological characteristics of IC neurons. **a**, Dendritic and local axonal morphologies of example IT, PT, and CT neurons from the aIC, mIC, and pIC. Local axons are defined as comprising the primary axonal trunk and its collateral branches extending from the soma, with termination points constrained to within 1 mm path distance from the parent cell body. Local axons, magenta; dendrites, cyan; somata, orange. **b**, Laminar density distributions of dendrites and local axons for IT, PT, and CT neurons. **c**, Left, radar plots of dendritic orientation relative to laminar direction for IT neurons in the aIC, mIC, and pIC. Right, quantification of dendritic angles (Kruskal–Wallis test followed by Dunn’s multiple comparisons). **d**,**e**, Same as **c** except for PT (**d**) and CT (**e**) neurons. **f**, Directional deviation of dendritic orientation for IT, PT, and CT neurons in the aIC, mIC, and pIC. A, anterior; P, posterior; S, superficial; D, deep. **g**, Left, axial, sagittal, and coronal views of first-order axonal collaterals from IT neurons, classified into five groups (color-coded) using the FNT-dist algorithm. Right, proportions of projection intensity from each collateral group to ipsilateral and contralateral brain regions. **h**,**i**, Same as **g** except for PT neurons (**h**) and CT neurons (**i**). **j**, Top, expansion rates quantifying the contribution of IT, PT, and CT collaterals to projections outside the IC (Kruskal–Wallis test followed by Dunn’s multiple comparisons). Bottom, expansion rates of collaterals from aIC, mIC, and pIC neurons (Kruskal– Wallis test followed by Dunn’s multiple comparisons). The expansion rate is defined as the ratio of the cumulative length of all axonal collaterals to the length of the primary axonal trunk. **k**, Expansion rates of collaterals for all 12 clusters (cluster 5 vs. 6, cluster 7 vs. 8: Kruskal–Wallis test followed by Dunn’s multiple comparisons; cluster 1-4 vs. 9-12: Wilcoxon rank sum test). Data are presented as mean ± s.e.m. n.s., P > 0.05; *P < 0.05; ****P < 0.0001.

**Extended Data Fig. 3.**
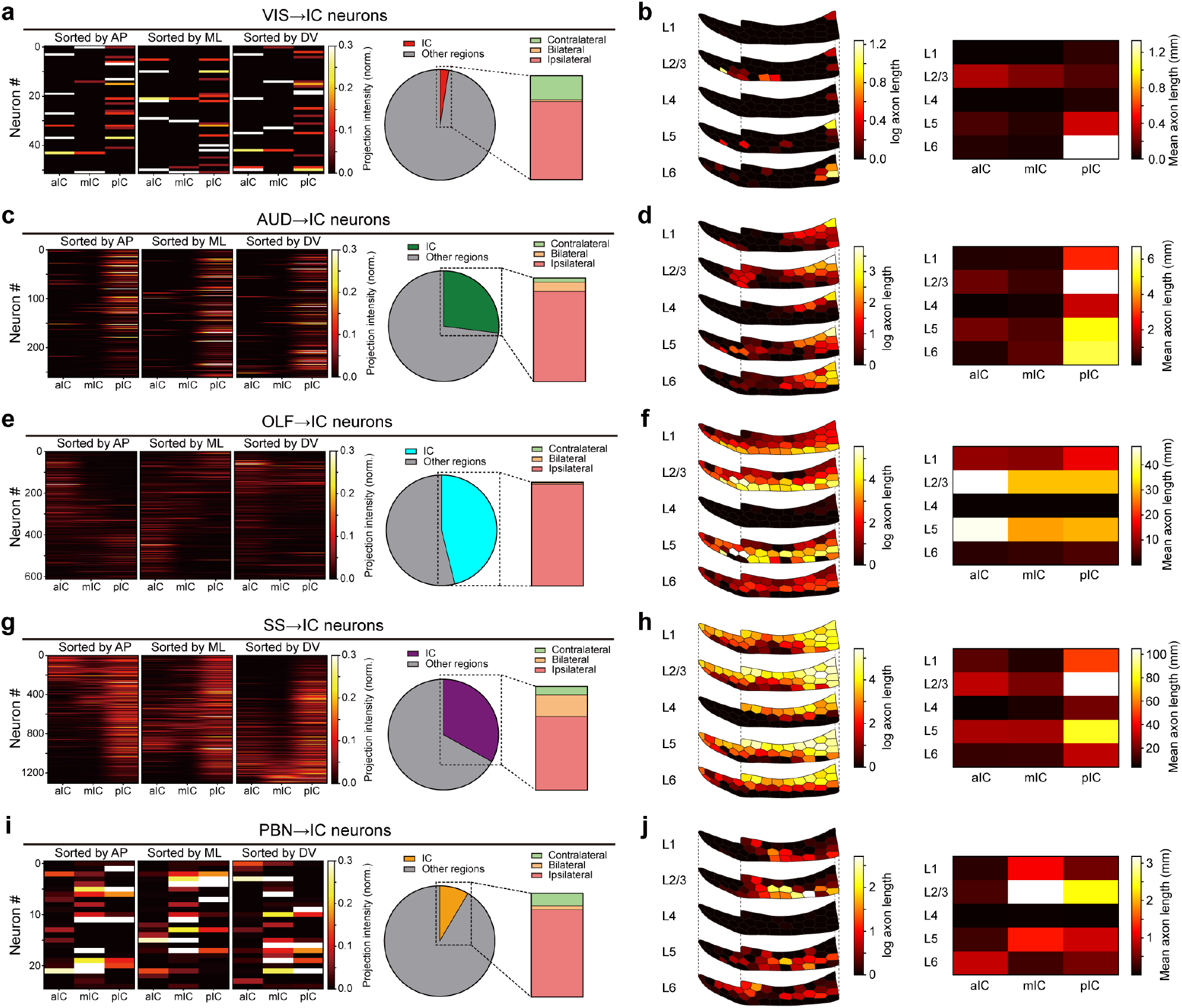
Topographic gradients and laminar preferences of sensory inputs to the IC. **a**, Left, normalized projection intensities from VIS-IC neurons to the aIC, mIC, and pIC, sorted by increasing values of somatic AP, ML, or DV coordinates. Right, proportions of VIS neurons projecting to the IC, subdivided into ipsilateral-only, contralateral-only, and bilateral projection patterns. **b**, Left, laminar topography of log-transformed projection intensities from VIS-IC neurons. Right, mean projection intensities to different layers. **c**,**e**,**g**,**i**, Same as **a** except for AUD-IC (c), OLF-IC (e), SS-IC (g), and PBN-IC (i) neurons. **d**,**f**,**h**,**j**, Same as **b** except for AUD-IC (d), OLF-IC (f), SS-IC (h), and PBN-IC (j) neurons.

**Extended Data Fig. 4.**
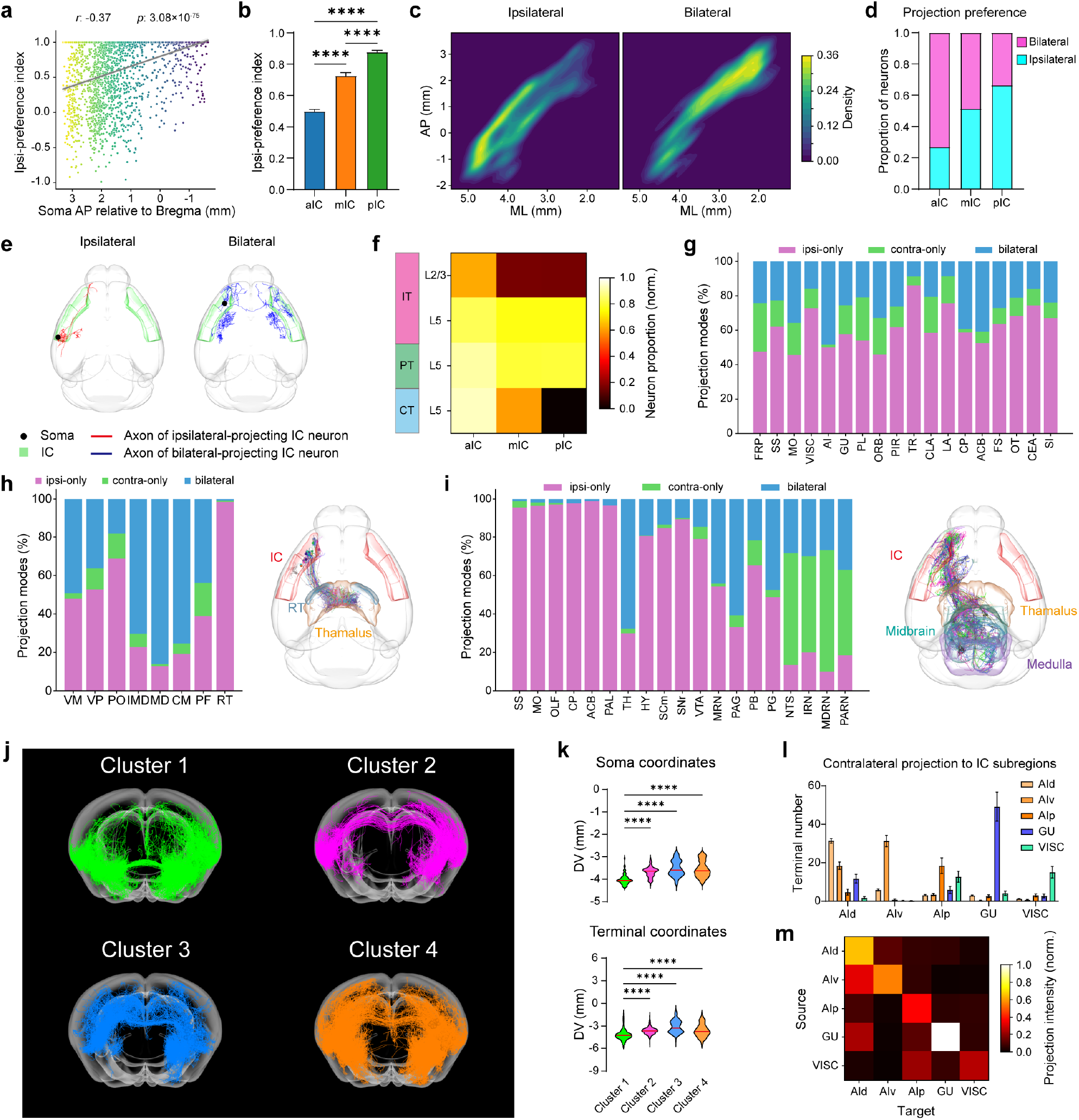
Interhemispheric projection patterns of IC neurons. **a**, Correlation between the ipsi-preference index of IC neurons and their somatic AP coordinates relative to Bregma. Color gradient indicates AP values of somata. Gray line denotes least-squares linear regression fit to the data. **b**, Ipsi-preference indices for neurons in the aIC, mIC, and pIC (Kruskal– Wallis test followed by Dunn’s multiple comparisons). **c**, Somatic density distributions of neurons exhibiting ipsi-only versus bilateral projections, color-coded by soma density. **d**, Proportions of ipsi-only and bilateral projecting neurons across the aIC, mIC, and pIC. **e**, Representative morphologies of IC neurons with ipsi-only versus bilateral projection patterns. **f**, Laminar distribution of bilateral-projecting IC neurons, showing enrichment in deep layers. **g**, Projection-mode proportions (ipsi-only, contra-only, bilateral) of IT neurons to their main targets. **h**, Left, projection-mode proportions of CT neurons to their major thalamic targets. Right, Projectomes of 40 randomly color-coded CT neurons. IC, thalamus, and reticular nucleus of thalamus are indicated in red, orange, and indigo, respectively. **i**, Left, projection-mode proportions of PT neurons to their main targets. Right, Projectomes of 20 randomly color-coded PT neurons. IC, thalamus, midbrain, and medulla are indicated in red, orange, green, and purple, respectively. **j**, Coronal views of projectomes from representative neurons in clusters 1–4. **k**, Top, somatic DV coordinates of neurons in clusters 1–4 (Kruskal–Wallis test followed by Dunn’s multiple comparisons). Bottom, DV coordinates of contralateral terminals for neurons in clusters 1–4 (Kruskal–Wallis test followed by Dunn’s multiple comparisons). **l**, Projection intensities from ipsilateral insular subregions to contralateral IC subregions. **m**, Heatmap showing IT neurons in a IC subregion prefers to innervate the contralateral homonymous subregion. Data are presented as mean ± s.e.m. **** P < 0.0001.

**Extended Data Fig. 5.**
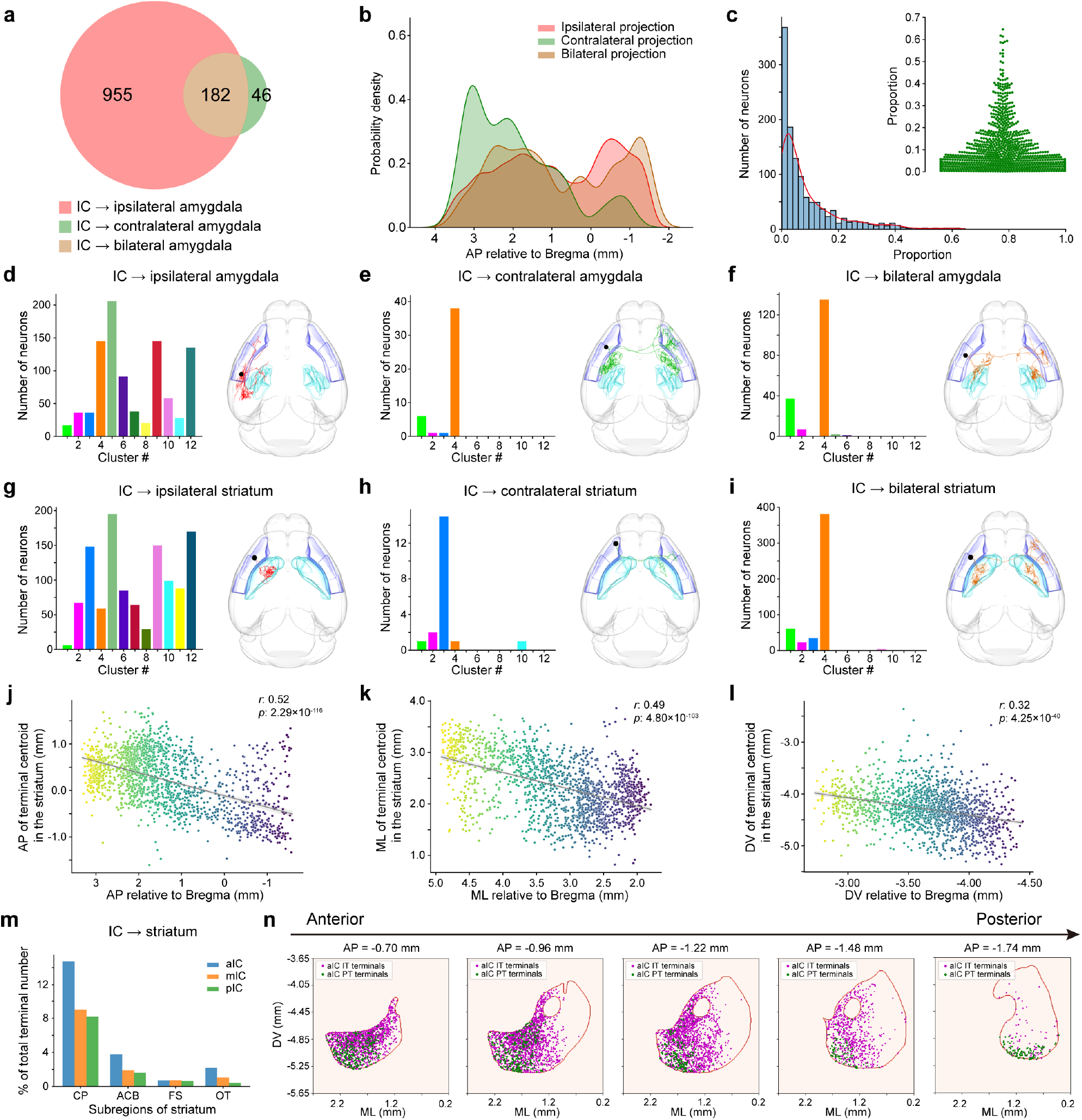
Distribution and projection patterns of IC neurons targeting the amygdala and striatum. **a**, Venn diagram showing the numbers of IC neurons projecting to the ipsilateral, contralateral, and bilateral amygdala. **b**, AP-axis probability density distributions of somata of IC neurons projecting to the ipsilateral, contralateral, or bilateral amygdala. **c**, Relationship between the proportion of amygdalar terminals relative to total terminals in amygdala-projecting IC neurons and the corresponding neuron counts. The red line indicates the kernel density estimation (KDE) fit. Inset, truncated swarmplot of the same data. **d-f**, Left, distribution of IC neurons projecting to the ipsilateral (**d**), contralateral (**e**), and bilateral (**f**) amygdala across the 12 clusters. Right, representative morphologies of IC neurons corresponding to each projection type. **g-i**, Left, distribution of IC neurons projecting to the ipsilateral (**g**), contralateral (**h**), and bilateral (**i**) striatum across all 12 clusters. Right, representative morphologies of IC neurons corresponding to each projection mode. **j-l**, Linear correlations between the centroid coordinates of striatal terminals and the somatic coordinates of IC neurons projecting to the striatum. Color gradient indicates AP/ML/DV values of somata. Gray line denotes least-squares linear regression fit to the data. **m**, Projection intensities from IC neurons to striatal subregions, expressed as the proportion of terminals within each subregion relative to the total terminals arising from each IC subdivision. **n**, Spatial heterogeneity of aIC-derived IT/PT neuron terminals in the nucleus accumbens (ACB) across multiple AP planes relative to the Bregma.

**Extended Data Fig. 6.**
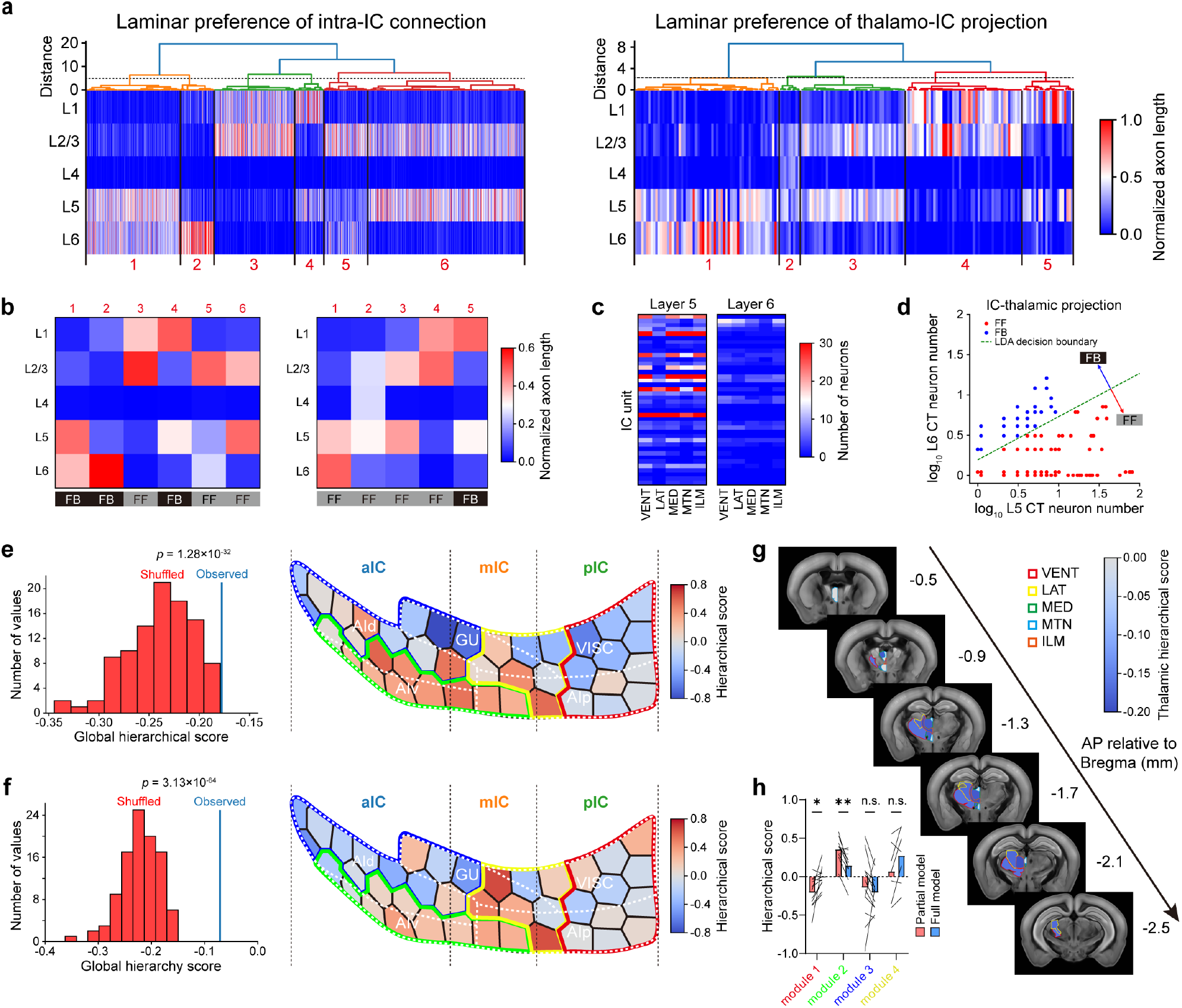
Hierarchy of IC-thalamus connectivity. **a**, Left, laminar preferences of intra-IC connections, revealing six characteristic patterns identified by hierarchical clustering. Right, laminar preference patterns of thalamo-IC projections from VENT, LAT, MED, MTN, and ILM to the 40 IC units (VENT: n = 119; LAT: n = 46; MED: n = 126; MTN: n = 110; ILM: n = 90). **b**, Classification of intra-IC (left) and thalamo-IC (right) feedforward (FF) and feedback (FB) projections based on laminar preferences identified in **a. c**, number of layer 5 and 6 CT neurons projecting to five thalamic subregions from each IC unit. **d**, FF/FB classification of IC-thalamic projections using linear discriminant analysis (LDA). **e**, Left, partial hierarchy model incorporating intra-IC connections only, showing a significantly higher global hierarchy score compared to shuffled controls (P = 1.28 × 10^−32^, one-sample t-test). Right, resulting IC hierarchy map, with hierarchy scores color-coded across IC units. Module boundaries are shown in distinct colors; white dashed lines denote five IC subregions. **f**, Same as **e** except for the full IC–thalamus hierarchy model (P = 3.13 × 10^−64^, one-sample t-test). **g**, Hierarchy map of five thalamic subregions in the full model, with region boundaries and hierarchy scores indicated. **h**, Hierarchy scores of each IC module for partial and full models (paired t-test for each module). n.s., P > 0.05; *P < 0.05; **P < 0.01.

**Extended Data Fig. 7.**
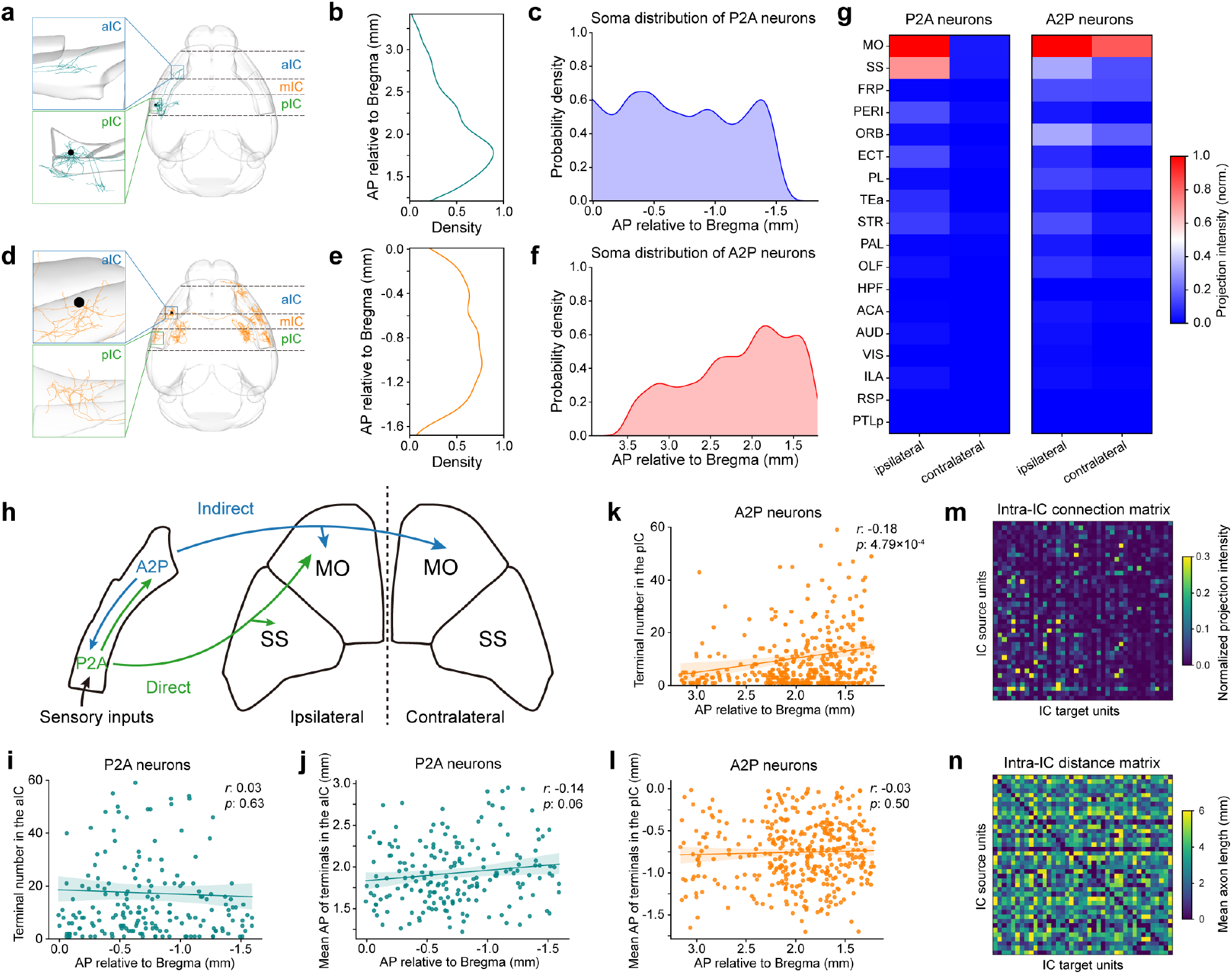
Intra-IC connectivity rules of IC neurons. **a**, Single-neuron morphology of an example P2A neuron projecting from the pIC to the aIC. Soma, black dot. **b**, AP-axis density distribution of P2A terminals within the aIC. **c**, AP-axis probability density distribution of P2A somata. **d-f**, Same as **a-c** except for A2P neurons. **g**, Normalized projection intensities to extra-IC brain regions for P2A (left) and A2P (right) neurons. **h**, Schematic of information flow from P2A and A2P neurons to motor (MO) and somatosensory (SS) cortices, highlighting their major projection pathways. **i**, Correlation between aIC terminal counts of individual P2A neurons and their somatic AP coordinates. **j**, Correlation between AP coordinates of P2A terminals and their somatic AP positions. **k**,**l** Same as **i**,**j** except for A2P neurons. **m**, Normalized intra-IC connection matrix for the rate-based whole-IC model. **n**, Mean axon lengths between IC units, estimating relative propagation delays for the rate-based whole-IC model.

**Extended Data Fig. 8.**
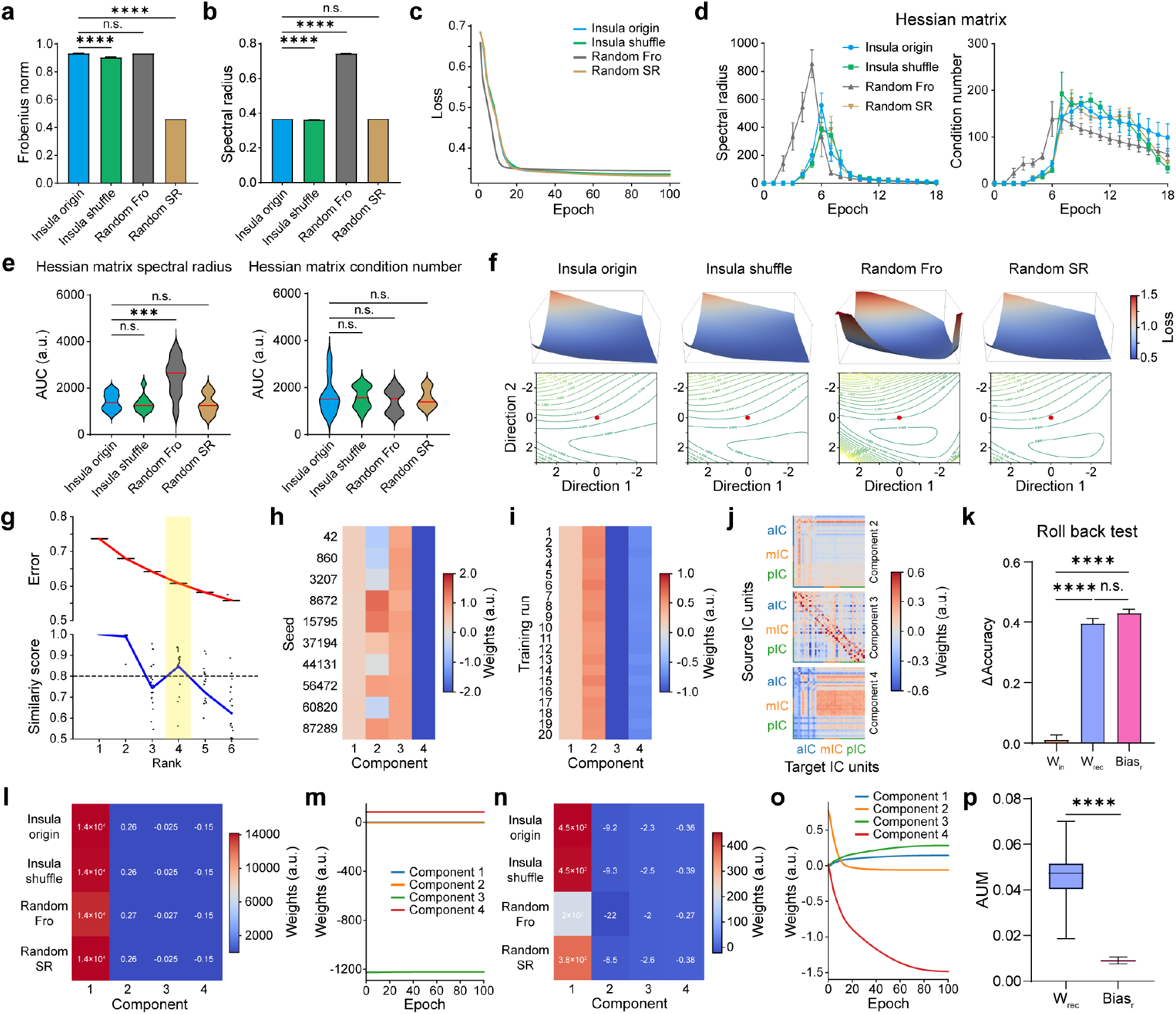
Topological priors, rather than basic matrix statistics, drive accelerated learning in IC-inspired RNNs. **a**, Comparison of Frobenius norms of recurrent weight matrices across RNN groups relative to the Insula-origin group (Kruskal-Wallis test followed by Dunn’s multiple comparisons test). **b**, Same as **a** except for spectral radii (Kruskal-Wallis test followed by Dunn’s multiple comparisons test). **c**, Training loss trajectories for the four RNN groups. The loss decreased rapidly during the first 18 epochs and subsequently plateaued, indicating fast initial convergence followed by slower optimization progress. Therefore, we define the first 18 epochs as the early-training phase. **d**, Early-training dynamics: spectral radius (ρ; left) and condition number (κ; right) of Hessian matrices across groups. **e**, Quantification of AUC for spectral radius and condition number during early training shown in **d**. (Kruskal-Wallis test followed by Dunn’s multiple comparisons test) **f**, Top, loss-landscape surfaces for initialized networks across groups. Bottom, corresponding 2D contour plots, with the red dot marking the origin. **g**, Top, model reconstruction error as a function of the number of components with each dot representing a single optimization run. Bottom, similarity score by model component. Each dot represents the similarity of an individual optimization run relative to the best-fit model in its category. The yellow rectancle shows the selected number of components. **h**, Seed-factor weights from fifth-order tensor decomposition of recurrent weight matrices. **i**, Run-factor weights from the same decomposition. **j**, Weight-factor components 2–4 derived from tensor decomposition of recurrent weight matrices. **k**, Rollback analysis showing test-set accuracy after resetting input weights, recurrent weights, or bias to their untrained states (Kruskal-Wallis test followed by Dunn’s multiple comparisons test). **l**, Method-factor weights for input-weight tensors across the four initialization schemes. **m**, Epoch-factor trajectories for input weights across training. **n**, Method-factor weights for bias tensors across initialization schemes. **o**, Epoch-factor trajectories for bias across training. **p**, Average update magnitudes (AUM) for recurrent weights and bias during early training (Wilcoxon rank sum test). Data are presented as mean ± s.e.m. n.s., P > 0.05; ***P < 0.001; ****P < 0.0001.

**Extended Data Fig. 9.**
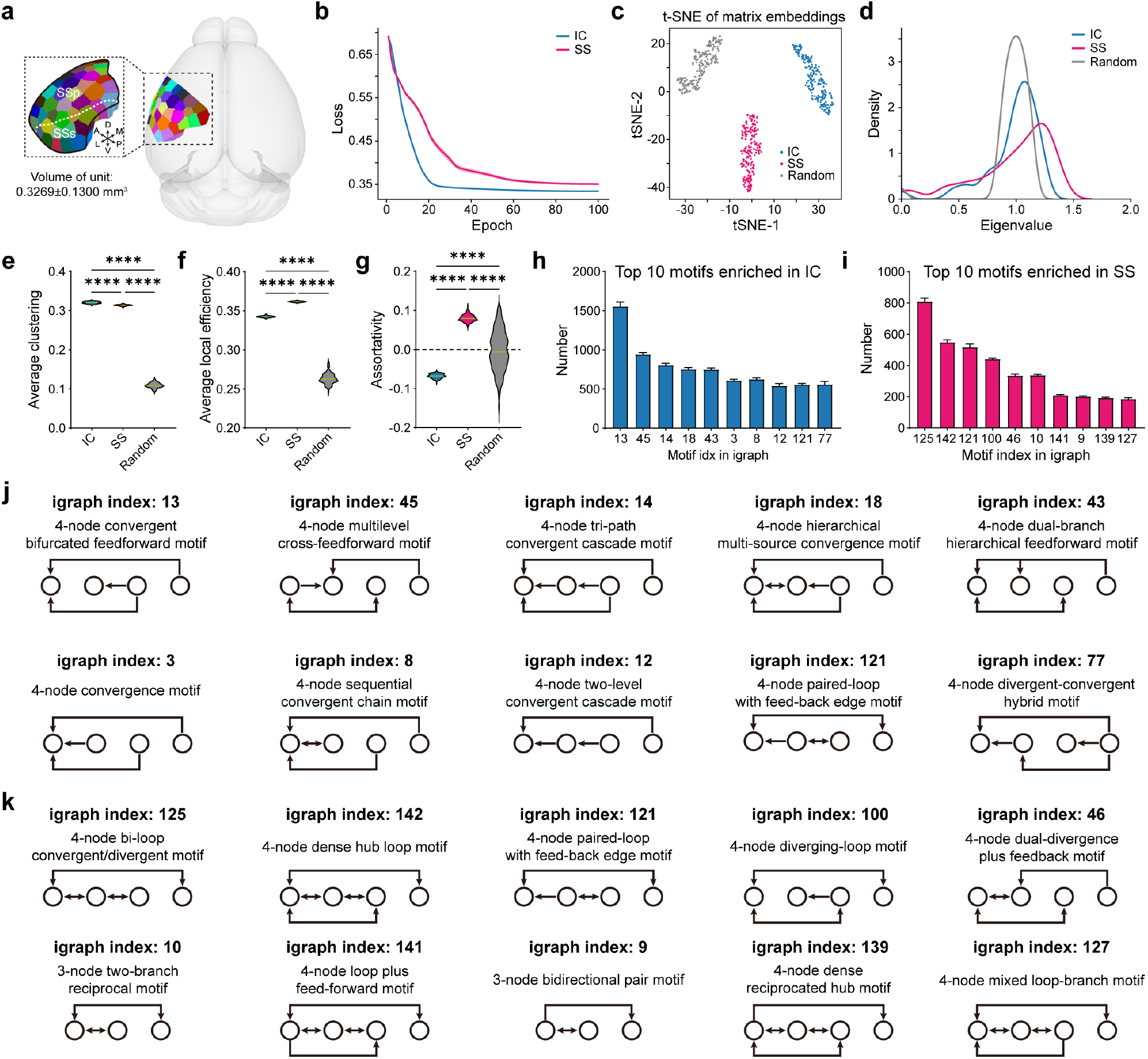
SS architecture exhibits topological features distinct from IC architecture. **a**, Schematic parcellation of the SS into 40 cortical units (mean ± s.d. volume: 0.3269 ± 0.1300 mm^3^), each randomly color-coded. **b**, Training loss trajectories for the IC and SS groups. **c**, t-SNE embedding of 32-dimensional GCN features derived from initialized recurrent weight matrices of IC-based, SS-based, and random RNNs; each point represents one matrix color-coded by network type. **d**, Spectral density distributions of normalized Laplacian eigenvalues for intra-IC, intra-SS, and random connectivity matrices. **e-g**, Network-metric comparisons across IC, SS, and random matrices, including average clustering coefficient (**e**), average local efficiency (**f**), and assortativity (**g**). ****P < 0.0001. Kruskal-Wallis test followed by Dunn’s multiple comparisons. **h**,**i** Top 10 motifs enriched in the IC (**h**) and SS (**i**) architectures, identified using igraph. **j**,**k** Schematic representations of the motifs listed in **h** and **i**, respectively.

**Extended Data Fig. 10.**
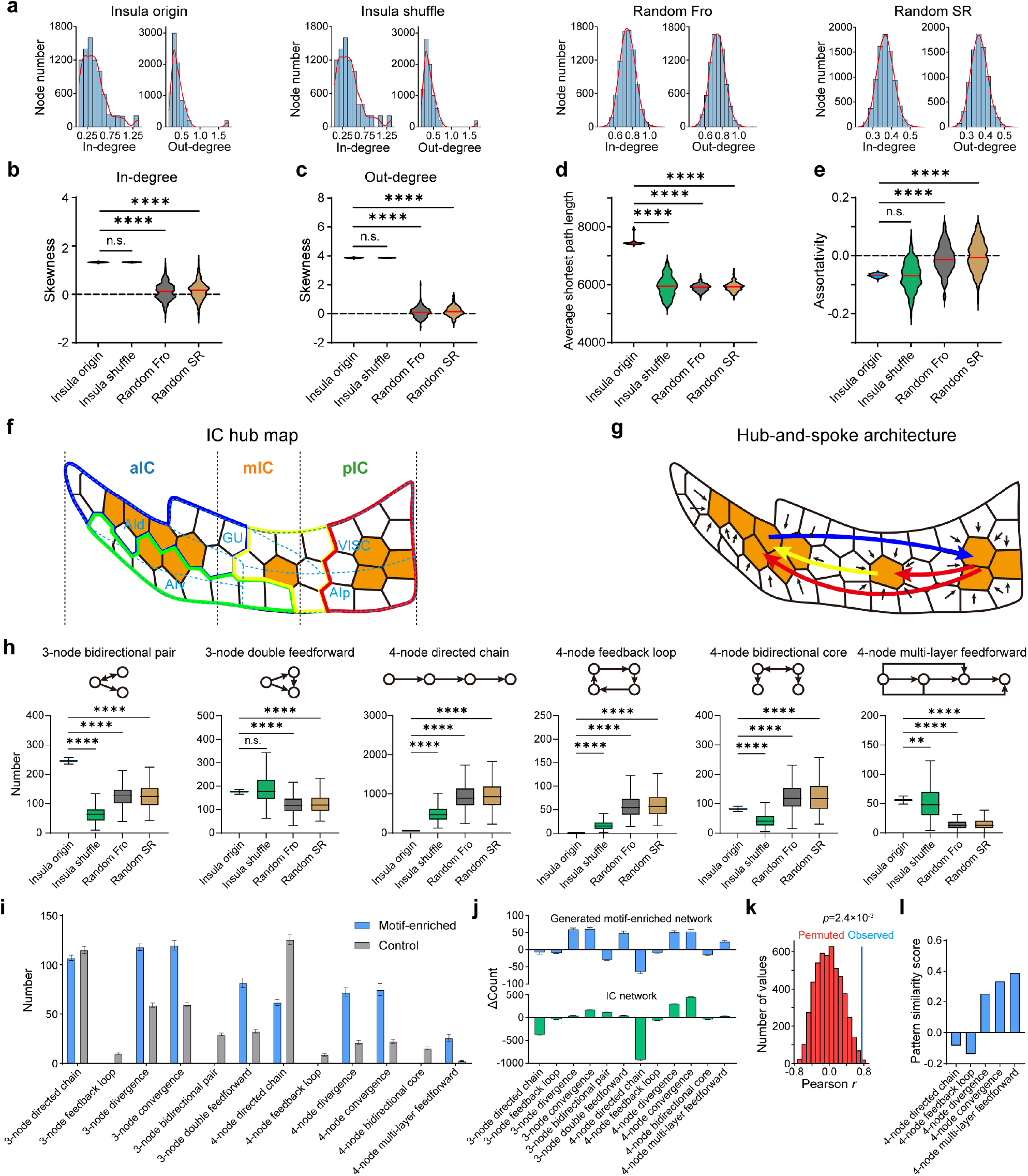
Topological features of IC architecture reveal hub organization and motif enrichment. **a**, Distributions of weighted in-degree and out-degree for the four recurrent weight matrix types (n = 200 per group), with red lines indicating KDE fits. **b**,**c** Skewness of weighted in-degree (**b**) and out-degree (**c**) distributions across the four matrix types. n.s., P > 0.05; ****P < 0.0001. Kruskal-Wallis test followed by Dunn’s multiple comparisons. **d**,**e** Comparisons of average shortest path length (**d**) and assortativity (**e**) across matrix types. n.s., P > 0.05; ****P < 0.0001. Kruskal-Wallis test followed by Dunn’s multiple comparisons. **f**, Spatial topography of hub nodes within the insula, with hubs shown in orange. Boundaries of modules 1–4 are outlined in red, green, blue, and yellow, respectively. IC subregions are indicated by dashed light-blue contours. **g**, Schematic of the hub-and-spoke organization of the insula, with hub nodes shown in orange. Red, yellow, and blue arrows denote predominant long-range information flows from hubs in modules 1, 4, and 3, respectively; small black arrows represent local information flow. **h**, Frequencies of six additional representative motifs across the four recurrent weight matrix types. ****P < 0.0001; Kruskal-Wallis test followed by Dunn’s multiple comparisons. **i**, Frequencies of selected motifs in networks generated by directional embedding of the 4-node multilayer feedforward motif, compared with their corresponding control networks. **j**, Comparison of motif-enrichment patterns between the multilayer feedforward–enriched networks and the insular network. ΔCount indicates the difference in motif frequency between each target network and its matched control. **k**, Permutation test of Pearson correlation for the motif-enrichment patterns in **j**. Red histogram shows the distribution of correlations obtained from shuffled motif-enrichment patterns; the blue line marks the observed correlation. **l**, Pattern-similarity scores (see Methods) quantifying the similarity between the motif-enrichment profile of the IC network and those of networks enriched for specific motifs. Data are presented as mean ± s.e.m.

## Notes

### Competing Interest Statement

The authors have declared no competing interest.

